# A common regulatory switch controls a suite of C4 traits in multiple cell types

**DOI:** 10.1101/2023.12.21.572850

**Authors:** Daniel Camo-Escobar, Carlos Alcalá-Gutiérrez, Ernesto Palafox-Figueroa, Bruno Guillotin, Marcela Hernández-Coronado, José L. Coyac-Rodríguez, Vincent E. Cerbantez-Bueno, Aarón Vélez-Ramírez, Stefan de Folter, Kenneth D. Birnbaum, Carlos Ortiz-Ramírez

**Affiliations:** UGA Laboratorio Nacional de Genómica para la Biodiversidad, CINVESTAV Irapuato, Guanajuato 36821, México; Center for Genomics and Systems Biology, Department of Biology, New York University, New York, NY 10003, USA; Laboratorio de Investigación Interdisciplinaria, ENES-León, Universidad Nacional Autónoma de México. Guanajuato 37684, México; Department of Botany and Plant Sciences, University of California, Riverside, CA, 92521, USA

## Abstract

The C4 photosynthetic pathway provided a major advantage to plants growing in hot, dry environments, including the ancestors of our most productive crops. Two traits were essential for the evolution of this pathway: increased vein density and the functionalization of bundle sheath cells for photosynthesis. Although GRAS transcriptional regulators, including SHORT ROOT (SHR), have been implicated in mediating leaf patterning in both C3 and C4 species, little is known about what controls the specialized features of the cells that mediate C4 metabolism and physiology. We show in the model monocot, *Setaria viridis*, that SHR regulates components of multiple cell identities, including chloroplast biogenesis and photosynthetic gene expression in bundle sheath cells, a central feature of C4 plants. Furthermore, we found that it also contributes to the two-cell compartmentalization of the characteristic four-carbon shuttle pathway. Disruption of SHR function clearly reduced photosynthetic capacity and seed yield in mutant plants under heat stress. Together, these results show how cell identities are remodeled by *SHR* to host the suite of traits characteristic of C4 regulation, which are a main engineering target in non-C4 crops to improve climate resilience.

## Main

C4 photosynthesis has evolved from a C3 ground state as a carbon concentration mechanism that greatly increases plant productivity in warm and dry environments. It relies on shuttling four-carbon molecules across two different cell types in the leaves, mesophyll (m) and bundle sheath (bs), with the purpose of concentrating CO_2_ in the latter^1^. Such a concentration mechanisms is possible because bundle sheath cells are suberized, preventing both CO_2_ and oxygen from diffusing freely^2^. Thus, during heat stress when stomata are closed, CO_2_ levels can be kept high while oxygen is excluded, greatly limiting photorespiration and creating the ideal conditions for carbon fixation by rubisco.

However, in the C3 ancestors of C4 plants bundle sheath cells do not express the genetic toolkit for carbon fixation and photosynthesis^3^. Hence, a major evolutionary step in the transition to a C4 pathway was the reprogramming of the bundle sheath for photosynthesis. Indeed, it has been observed that chloroplast biogenesis and activation of photosynthetic functions in this cell type are essential steps in the transition from all C3 to C4 intermediates^4^. Moreover, this is the only feature that is universally shared between all C4 species, highlighting its importance^5^.

Because the bundle sheath becomes the main site of photosynthesis in C4 plants, as opposed to the more abundant mesophyll in C3, many species have evolved anatomical adaptations to increase bundle sheath to mesophyll ratio. One common adaptation is to increase vein density, which leads to an increment in the associated bundle sheath tissue^6^. Higher density is achieved through the development of more lower rank veins. Vein classification depends on their ontogeny: in grasses, mid and lateral veins are the first to develop, while rank1(r1) veins are formed after. Only C4 plants can produce additional files of smaller veins denominated rank2(r2) veins^4^.

There is evidence that members of the *SHORT ROOT* (*SHR*)*/SCARECROW* (*SCR*) pathway are involved in leaf patterning. *SCARECOW* mutants in maize show defects in mesophyll development that alter vein density by decreasing the number of cells between adjacent veins^7^. On the other hand, *SHORT ROOT* maize mutants show a subtle increase in the number of merged veins^8^, and it was recently demonstrated that mutants in rice also have mesophyll proliferation defects^9^. However, their role in the formation of the C4-specific r2 veins has not been determined.

Information is far more limited in the case of bundle sheath fate specification and functionalization for photosynthesis. In *Arabidopsis thaliana* -a C3 model-SHR has also been implicated in bundle sheath formation, but this has not been confirmed at the molecular level. Furthermore, the maize *GOLDEN2 (G2)* gene is the only known regulator involved in bundle sheath chloroplast biogenesis^4^. Nevertheless, this function is not always conserved across species, and although *g2* mutants have defects in chloroplast morphology, they do not show differences in the compartmentalized expression of the carbon shuttle enzymes that maintain the C4 cycle^10^. Therefore, regulators controlling this trait have not been described.

### *SHR* controls C4 vein development

The *SHR/SCR* pathway has been implicated in patterning roots and shoots. Hence, we first determined if *SHR* has a role in the development of *Setaria* C4 leaves. Two paralogs have been reported in this species: *SvSHR1* and *SvSHR2*. We evaluated the phenotype of CRISPR-Cas9 loss-of-function mutants for both genes, as well as a double mutant line^11^. *Svshr1* mutant plants showed a slight but significant reduction in both plant height and leaf width, while *Svshr2* mutants didńt have any obvious growth defects. *Svshr1/2* double mutant plants were severely stunted and developed very small and thin leaves (Fig. 1, a, b, e-g). These plants didńt grow past 3-4 weeks and never set any seeds.

**Fig. 1.**
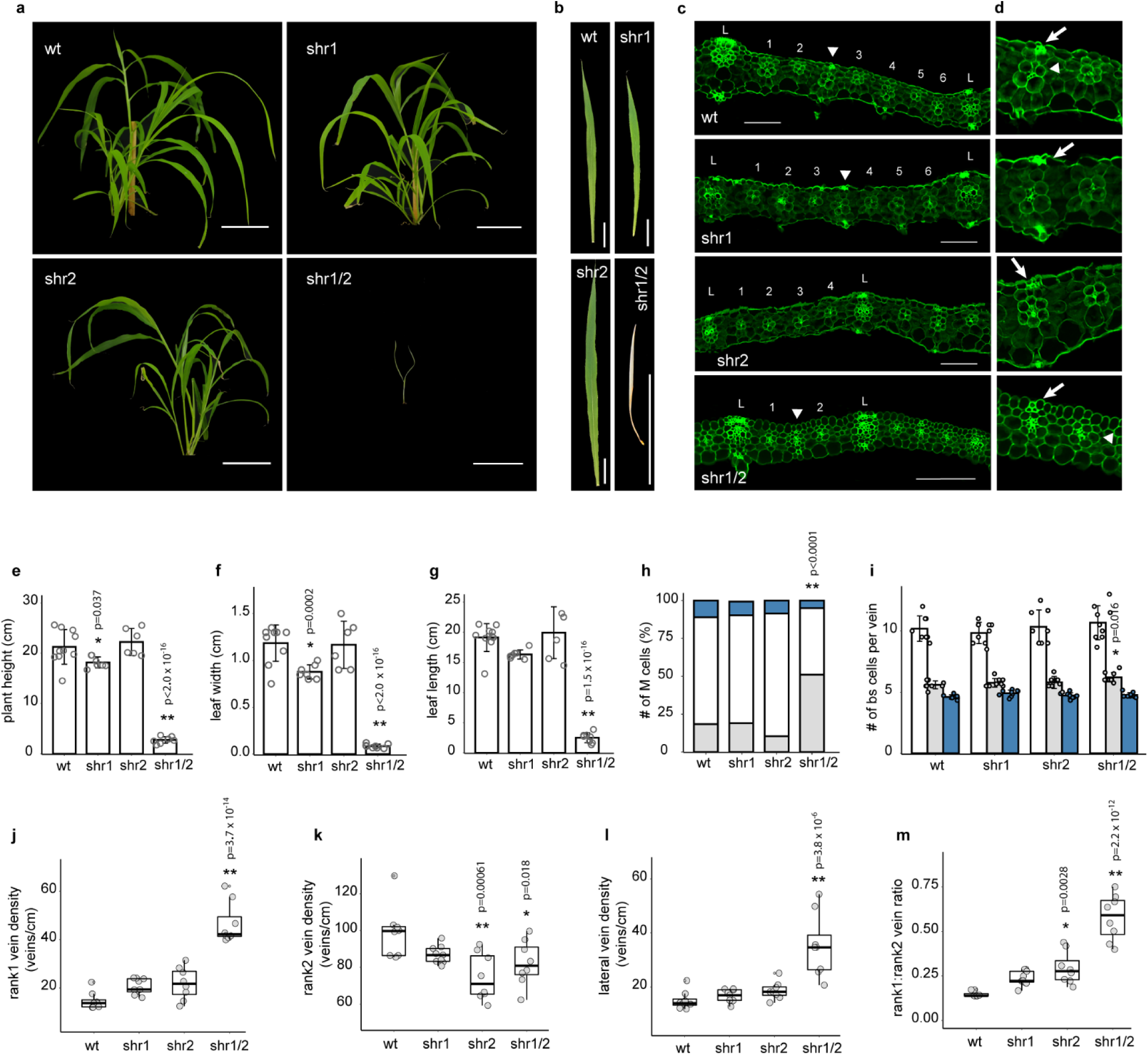
Anatomical and cellular phenotypic analysis of SHR mutants. (**a**) Representative images of wt, *Svshr1* single, *Svshr2* single, and *Svshr1/2* double mutant plants. (**b**) Representative images of the fourth emerged leaf of wt, *Svshr1* single, *Svshr2* single, and *Svshr1/2* double mutants. (**c**) Histological cross sections of mature Setaria wt, *Svshr1* single, *Svshr2* single, and *Svshr1/2* double mutant leaves showing lateral (L), rank1 (arrowheads), and rank2 veins (numbers). (**d**) Enlarged regions from (c) showing sclerenchyma (arrows) and bundle sheath (arrow heads). (**e**-**f**) Measurements of plant height, leaf width and length for Setaria wt (*n* =10), *Svshr1* (*n* = 7) single, *Svshr2* (*n* =7) single, and *Svshr1/2* (*n* =8) double mutant plants. (**h**) Percentage of the number of mesophyll cells separating veins for each genotype (one cell = blue, two cells = white, three cells = grey). (**i**) Average number of bundle sheath cells associated to each vein type (laterals = white, rank1 = grey, rank2 = blue) for *Setaria* wt (*n* =8), *Svshr1* (*n* = 9) single, *Svshr2* (*n* =8) single, and *Svshr1/2* (*n* =9) double mutant plants. (**j**-**l**) Vein type density quantification reported as veins/cm for wt (*n* =8), *Svshr1* (*n* = 9) single, *Svshr2* (*n* =8) single, and *Svshr1/2* (*n* =8) double mutant plants. (**m**) Quantification of the ratio of rank1 to rank2 veins for all genotypes (*n* as in j-l). Individual data points are shown as either open or closed grey circles. Asterisks indicate statistically significant difference with respect to wt (* = p < 0.05, ** = p < 0.001, Dunnet test after one way analysis of variance on all genotypes). Scale bars, 4 cm (a), 2 cm (b), and 100 µm (c). Box plot center lines represent the median and box limits indicate the 25^th^ and 75^th^ percentiles. Bar plots show the standard deviation of the mean.

Leaf sections were prepared to assess defects in patterning, particularly in the formation of r2 veins, which are specific to C4 plants and the most abundant vein type in these species. In wild-type plants, usually six r2 veins develop between each lateral vein (Fig. 1c, upper panel). The same pattern was present in *Svshr1,* which did not showed differences in any of the anatomical traits evaluated. In contrast, we observed that in *Svshr2* and *Svshr1/2* mutants less r2 veins were formed, only four and two veins between each lateral respectively (Fig. 1c, lower panels). To evaluate if patterning defects had an impact on vein density, we quantified vein types per cm. Again, both *Svshr2* and *Svshr1/2* showed reduced r2 vein density, 74 and 82 veins/cm respectively compared to 100 in wild type (Fig. 1k).

Notably, r1 and lateral veins were not affected in *Svshr2,* suggesting that the *SHR2* homolog specifically regulates r2 vein proliferation (Fig. 1j & l). The same was not observed in the double mutant, which showed an increased r1 and lateral vein density. We hypothesized this increment was an indirect consequence of impaired mesophyll cell division, leading to closer veins. Indeed, both SCR and SHR have been associated with this function^7,9^. We quantified the number of mesophyll cells separating each vein and as expected, the percentage of veins separated by just one cell instead of two was higher in *Svshr1/2*, 51% compared to 18% in wild type (Fig. 1h). Ultimately, these phenotypes resulted in a distorted proportion of vein types as evidenced by the high rank 1 to rank 2 ratios in both mutants (Fig. 2m). Taken together, our results suggest that *SHR* promotes development of rank 2 veins, which are unique to Kranz anatomy, and that it can modulate r1 and lateral vein density through mesophyll proliferation.

**Fig. 2.**
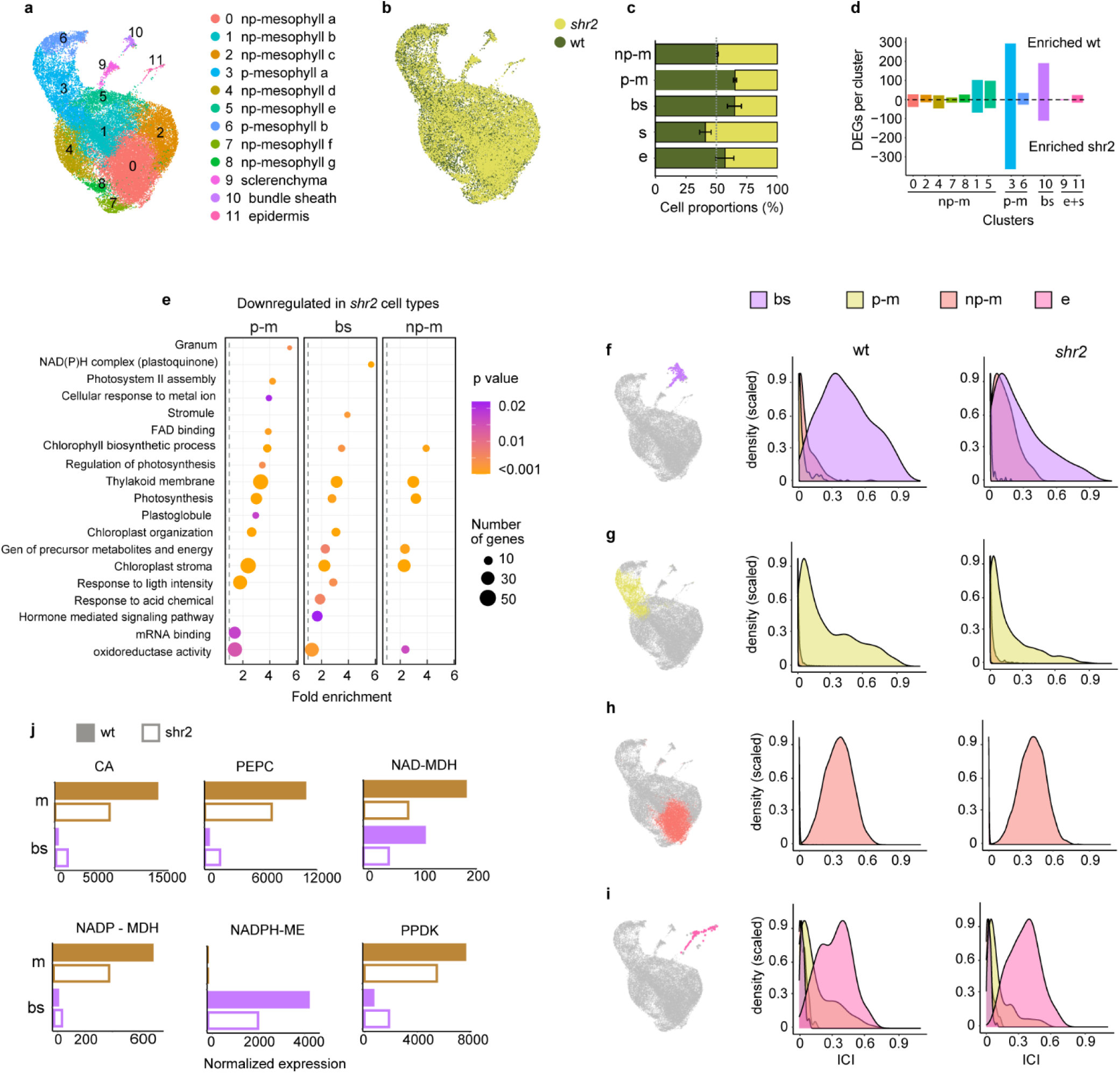
Single-cell RNA-seq and quantification of cellular identities. (**a**) UMAP representation of single-cell transcriptomes showing cluster identities as determined by expression of canonical tissue markers (photo = photosynthetic). (**b**) Integrated (batch corrected) single cell transcriptomes of wt and Svshr2 genotypes mapped onto the same UMAP depicted in (a). (**c**) Percentage of wt and Svshr2 cells across different cell types (M, mesophyll; MP, photosynthetic mesophyll; BS, bundle sheath; S, sclerenchyma; E, epidermis). Error bars represent 95% binomial confidence intervals. (**d**) Histogram of the number of differentially expressed genes (DEGs) between wt and Svshr2 across cellular clusters (more than 2-fold difference in expression). Cell type codes as in (c). (**e**) Gene ontology enrichment analysis of downregulated DEGs in Svshr2 across photosynthetic mesophyll (MP), bundle sheath (BS), and mesophyll (M). DEGs from clusters with the same identity were combined. No gene ontology category was enriched in other cell types. (**f**-**i**) Quantification of cellular states with the ICI method, showing cellular identity scores as density plots. Differences between wt and Svshr2 distributions were evaluated by measuring their empirical cumulative distribution function through a Kolmogorov-Smirnov test. Significant differences (p < 0.01) were found for (f) and (g). For visualization, analyzed cell populations are mapped onto the same UMAP depicted in (a) and highlighted in different colors (blue, bundle sheath; yellow, photosynthetic mesophyll; brown, mesophyll; pink, epidermis). (**j**) Expression differences of the main C4 cycle enzymes between wt (solid bars) and Svshr2 (outline bars) in two different tissues: bundle sheath (BS) mesophyll (M).

### Specification of photosynthetic function

In addition to patterning defects, mutant plants showed novel phenotypes at the cellular level: although cells at the bundle sheath position were present in comparable numbers (Fig. 1i), they lacked their characteristic morphology. In wild type, the bundle sheath was clearly distinguishable from mesophyll cells by its wreath arrangement and ticker cell walls that emit more autofluorescence (Fig. 1d, upper panel). However, in *Svshr1/2* leaves this distinction is no longer visible (Fig. 1d, lower panel). Indeed, suberin staining of mutant bundle sheath revealed a total absence of cell wall suberisation (Extended Data Fig.1), suggesting cell fate specification defects.

To explore how extensively SHR controls the anatomy and gene expression reprogramming associated with C4 photosynthesis, we perform a detailed comparison of gene expression patterns between wild type and mutant leaves using single-cell RNAseq. A protocol to generate *Setaria* leaf protoplast was optimized based on previous reports^11,12^. Protoplasts were generated from middle sections of fully expanded leaves and used to generate single-cell cDNA libraries with the 10x Genomics platform. We did not profiled *Svshr1* because of its lack of phenotype. Regarding *Svshr1/2,* protoplasts were very fragile and bursted after enzymatic digestion preventing single-cell profiling.

We generated a total of 18,352 and 15,994 high quality single cell transcriptomes from wild-type and *Svshr2* plants respectively, sequenced in three different batches. Wild-type cells were assigned to 12 clusters according to their transcriptomés global similarity and visualized in two dimensions using the uniform manifold approximation and projection (UMAP) method (Fig. 2a). We then determined cluster identity using canonical tissue-specific markers. For example, *NADPH-ME* and *RBCS1*, for bundle sheath, *CA*, *PDK1*, *MDH6* for mesophyll, and *GRP6* for epidermis. We independently confirmed the identity of relevant clusters using RNA-FISH (Extended Data Fig. 2).

As found in previous leaf single-cell studies in monocots, mesophyll cells were by far the most abundant ^13^. They grouped into a supercluster composed of 9 subclusters (Fig. 2a). To explore differences between mesophyll sub-populations, we identified differentially expressed genes across subclusters and performed gene ontology analyses. A clear pattern emerged: mesophyll clusters could either be classified as enriched in photosynthetic functions or enriched in non-photosynthetic functions (Supp. Table 1). Therefore, we classified clusters as photosynthetic mesophyll (p-m) and non-photosynthetic mesophyll (np-m) respectively. We also detected bundle sheath, epidermis and sclerenchyma cells, which formed smaller, very distinct clusters, separated from the mesophyll supercluster (Fig. 2a).

To perform comparative analyses across genotypes, we integrated wild-type and *Svshr2* mutant samples using “matched” transcriptomes as anchors with the CCA Seurat package (Fig. 2b). Once integrated, we first determined if cell-type proportions in leaves were changing due to the absence of *SHR*, as this could hint to cell-type specific developmental defects. We discovered that the proportion of both bundle sheath and photosynthetic mesophyll decreased significantly in *Svshr2* mutant leaves (Fig. 2c). Although cellular distribution can be affected by cell wall composition during protoplasting, this result likely reflects proliferation and/or differentiation defects on these cell types.

Notably, further comparative transcriptomic analyses suggested that SHR controls gene expression in bundle sheath cells that contributed to critical components of the C4 physiology. Bundle sheath and photosynthetic mesophyll cells had the highest number of differentially expressed genes (DEGs) between wild type and *Svshr2*, whereas other cell types showed only minimal changes to their transcriptomes (Fig. 2d). Moreover, gene ontology analyses on those DEGs showed that most genes affected in mutant bundle sheath were involved in photosynthetic functions and chloroplast biogenesis (Fig. 2e, Supp. Table 2). Strikingly, 16 out of 19 GO categories were related to photosynthesis (e.g. photosystem assembly, chloroplast biogenesis, chloroplast organization and photosynthetic regulation). This overrepresentation of photosynthesis related GOs was cell-type specific, because DEGs for other cell types like non-photosynthetic mesophyll only showed moderate enrichment, or in the case of epidermis and sclerenchyma, no enrichment in any GO category (Fig. 2e). Thus, in *Setaria*, SHR appears to be highly specific in imparting photosynthetic capacity to bundle sheath cells and photosynthetic mesophyll.

### SHR establishes bundle sheath identity

As mentioned before, two characteristics that define bundle sheath cells in C4 plants are the suberisation of its cell wall and their photosynthetic functionalization. Because both identity-defining aspects seem to be compromised in mutant plants, we proposed that *SHR* might be important in the establishment and maintenance of bundle sheath identity. To explore this possibility, we tracked cell identity changes in mutant plants using the index of cell identity (ICI). In brief, this algorithm can identify stable and transitional cell fates using large identity marker references^14^. In this case, we defined at least 100 marker gene sets for each cell type from our single cell data, and generated ICI scores for all wt and mutant cells.

We individually evaluated cells from four different clusters representing the most relevant types (e.g. bundle sheath, photosynthetic mesophyll, non-photosynthetic mesophyll, and epidermis), and visualized cell identity contributions as density plots. Notably, *Svshr2* bundle sheath cells showed the greatest loss of identity compared to wild type (Fig. 2f). Moreover, loss of bundle sheath fate was accompanied by a significant gain in non-photosynthetic mesophyll identity, resembling a transitional fate state characterized by cells with mixed identities. Importantly, this indicates that SHR not only promotes bundle sheath fate, but also represses non-photosynthetic mesophyll identity within these cells.

We further traced ICI scores in photosynthetic mesophyll and observed a similar trend, a significant loss of identity compared to wild type (Fig. 2g). In contrast, when we evaluated other cell populations like epidermis and non-photosynthetic mesophyll, there was no effect on cell identity due to *SHR* loss of function (Fig. 2, h & i). Taken together, these results suggest that *SHR* is important for fate specification of cells with photosynthetic functions. Notably, by promoting the expression of photosynthesis related genes in bundle sheath cells and simultaneously repressing markers of non-photosynthetic tissues, *SHR* seems to facilitate the functionalization of this tissue for photosynthesis.

### Downregulation of the C4 shuttle system

The C4 cycle depends on a series of compartmentalized enzymatic reactions that work by shuttling carbon molecules from mesophyll into bundle sheath cells. Thus, we examined downregulated genes in *Svshr2* mutants to assess if enzymes specific to the C4 pathway were affected. Five enzymes perform the main enzymatic steps: first, carbonic anhydrase (CA) catalyzes a primary CO_2_ fixation in the mesophyll by converting CO_2_ into HCO^-^. This is later used by phosphoenolpyruvate carboxylase (PEPC) and malate dehydrogenase (NADP-MDH) to generate the four carbon molecules that are translocated into the bundle sheath. Once there, CO_2_ is released from the four-carbon chains by NADP-malic acid (NADP-ME). Finally, the resulting C3 acids are shuttled back to mesophyll where PPDK uses them to regenerate PEP. Thus, in C4 plants these enzymes are highly compartmentalized, NADP-ME being bundle sheath specific, whilst the rest are mesophyll specific.

We quantified the expression of C4 shuttle enzymes in both cell types. As expected, mesophyll-associated enzymes were highly compartmentalized in wild type: *CA, PEPC, NADP-MDH and PPDK* were expressed at high levels in mesophyll cells and almost completely absent in bundle sheath (Fig. 2j). However, when *Svshr2* cells were analyzed, we found a marked reduction in mesophyll transcript levels (half compared to wild type) and a moderate increase of expression in bundle sheath cells, particularly for *PEPC, CA* and *PPDK*. Hence, expression specificity clearly decreased in mutant cells.

Transcript levels of the bundle sheath-associated enzyme *NADP-ME* were also reduced by half in *Svshr2*. However, in this case expression specificity was not affected as transcripts were absent from mesophyll cells. (Fig. 2j). Altogether, these results suggest that *SHR* directly or indirectly regulates the expression of the main carbon shuttle enzymes, contributing to their compartmentalization.

### *SHR* transcripts localize to bs cells

As shown before*, SHR* regulates gene expression in bundle sheath and photosynthetic mesophyll and is important for maintaining their identity. If this transcription factor has a cell-autonomous function, we hypothesized that transcripts should be localized to these tissues. To gain insight into the spatial pattern of *SHR* transcript accumulation, we used hybridization chain reaction (HCR) RNA-FISH technology. Specific probe sets were designed for *SvSHR1* and *SvSHR2* genes and half mount hybridization was performed on middle sections of partially emerged leaves^15^. Transcripts of both genes were mainly detected in bundle sheath cells associated with rank1 and rank2 veins but not in bundle sheath associated with mid and lateral veins, showing that expression in bundle sheath cells is specific depending on the type of vein (Fig. 3, a & b upper panel). Signal was also detected at low levels in the mesophyll cells directly in contact with bundle sheath, especially in the case of *SvSHR1* transcripts. Notably, *SvSHR2* transcript was also detected in the vasculature of lateral and mid veins (Fig. 3b, lower panel). Hence, the localization of *SHR* transcripts to the bundle sheath is probably important for driving the expression of photosynthetic genes in these cells.

**Fig.3.**
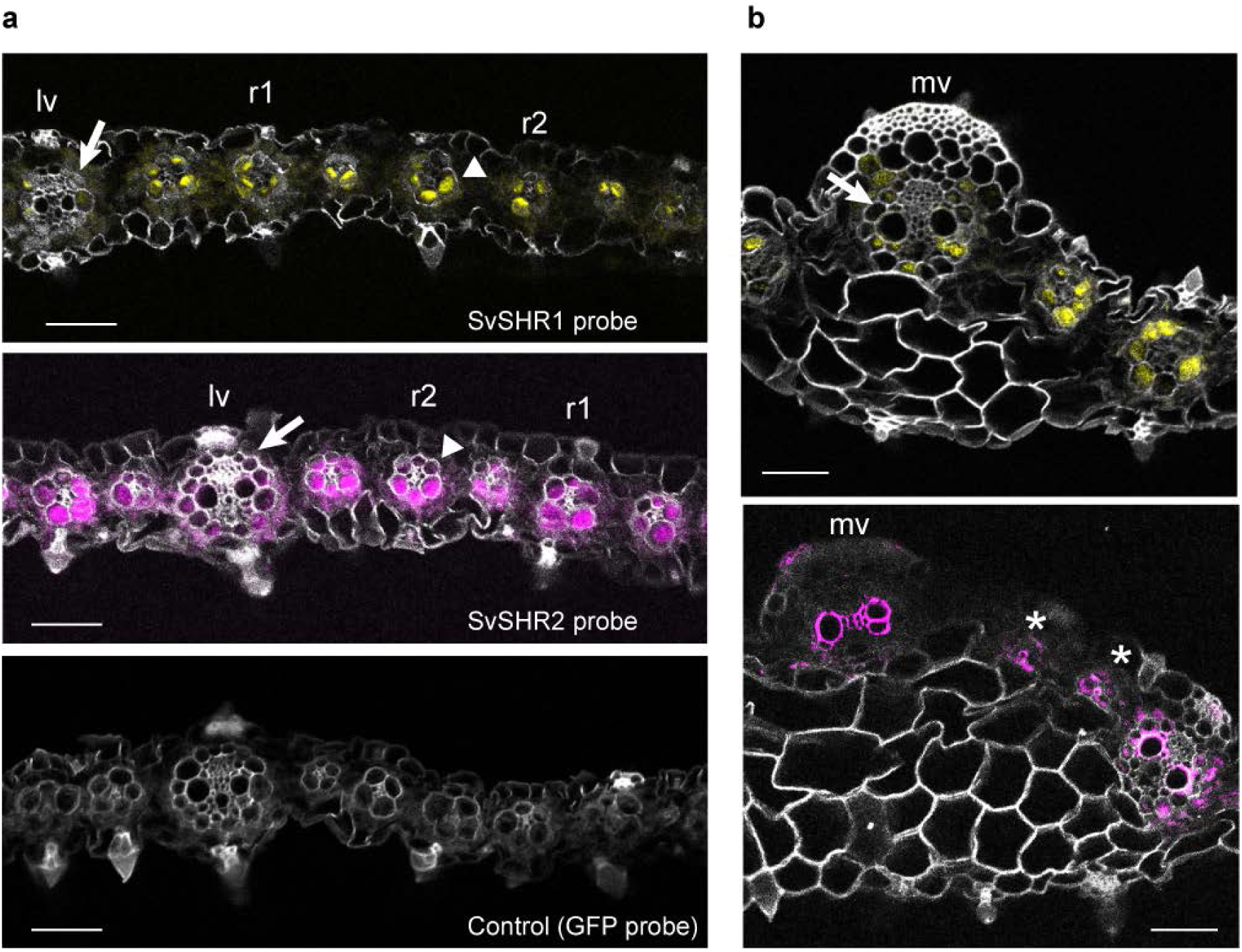
SHR transcript localization in *Setaria* leaves. (**a**) In situ hybridization (RNA FISH) showing the localization of *SvSHR1* (top), *SvSHR2* (middle), and a negative control probe (bottom). Signal was mainly detected in the bundle sheath of rank1 (r1) and rank2 (r2) veins (arrow heads), but not in lateral veins (lv, arrows). (**b**) In situ hybridization performed in the middle vein region of the leaf. *SvSHR1* signal (top) is also low on the bundle sheath of middle veins (bottom) and absent from parenchyma cells. *SvSHR2* transcripts are localized to the vascular tissue of middle veins (mv) and emerging small veins (asterisks). Yellow and purple correspond to Sv*SHR1* and *SvSHR2* signal respectively. Grey signal corresponds to cell wall autofluorescence. Scale bars, 50 µm.

### Cell fate defects impair photosynthesis

Given that in C4 plants the bundle sheath is the main site of photosynthesis and that mutant plants showed a loss of bundle sheath identity, we asked if this could impact photosynthetic rates. CO_2_ assimilation, A, versus substomatal CO_2_ concentration, Ci, curves were generated for wt, single and double mutants. Both *Svshr1* and *Svshr2* single mutants showed reduced photosynthetic rates at most substomatal CO_2_ concentrations measured, including at saturating CO_2_ (A_max_, Fig. 4, a & e). However, *Svshr2* plants clearly showed a more pronounced reduction (Fig. 4e). When measurements were done under reduced oxygen levels (2%), assimilation rates were more variable. This resulted in differences not being significant between wild type and mutant, especially for *Svshr1*, indicating photorespiration might influence assimilation rates differently in wild type and mutants (Fig. 4b). In the case of *Svshr1/2* double mutant, the phenotype was striking, since we could only detect negligible CO_2_ assimilation rates at saturating CO_2_ concentrations (1000 µmol mol^-1^) (Extended Data Fig.3). Therefore, double mutant plants cannot perform photosynthesis under normal conditions.

**Fig. 4.**
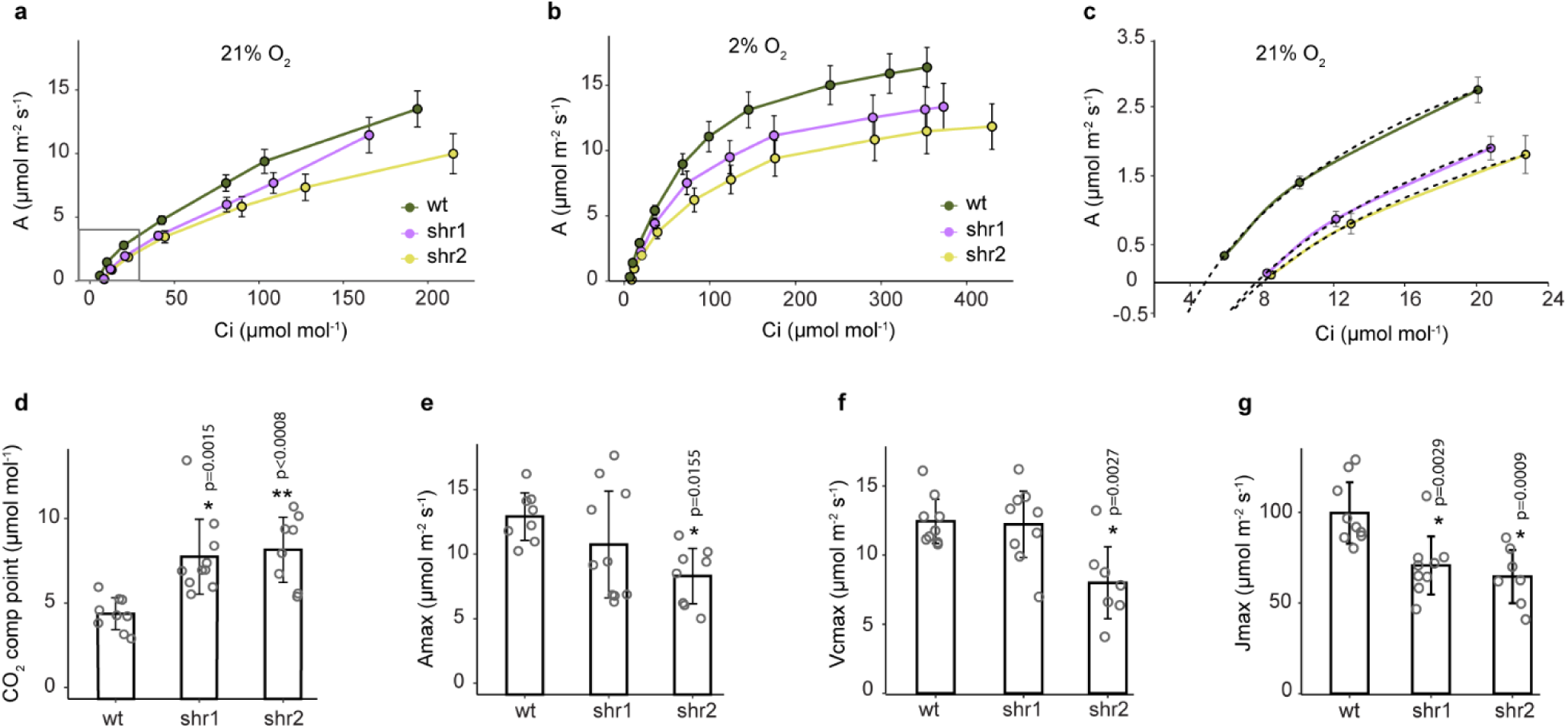
Changes in the photosynthetic rate of wt and SHR mutants. (**a**) A/Ci curves of wt (*n* = 9), shr1 (*n* = 10), and shr2 single mutants (*n* = 8) measured at 21% oxygen. Error bars are standard errors of the mean. (**b**) As in (a) except measurements were performed at 2% oxygen. (**c**) Enlarged region from grey box in (a) to show the shift in the CO2 compensation point between wt and mutants. Dash lines represents a polynomial regression fit on the A/Ci curves. (**d**) CO2 compensation. (**e**) Amax. (**f**) Vcmax. (**g**) Jmax. Individual data points are shown as open grey circles. Asterisks indicate statistically significant difference with respect to wt (* = p < 0.05, ** = p < 0.001, Dunnet test after one way analysis of variance on all genotypes). Bar plots show the standard deviation of the mean.

A defining feature of plants with a functional C4 cycle is a low CO_2_ compensation point ^16^. This is because C4 plants can actively concentrate CO_2_ in the bundle sheath, which enables positive photosynthesis rates at much lower atmospheric CO_2_ concentrations. To determine changes in the compensation point between wild type and mutants, we used a polynomial regression fit on A/Ci curves (Fig. 4c). Wt plants had a mean compensation point of 4.3 µmol/mol, which is in line with previous measurements for *Setaria*^17^. Remarkably, the CO_2_ compensation point of *Svshr1* and *Svshr2* mutants was almost twice as high (8 µmol/mol, Fig. 4d). Although this value is still far from that of C3 plants, it is an indication that the C4 cycle is not operating normally.

We further modelled A/Ci curves (von Caemmerer) to estimate two key parameters of photosynthetic kinetic processes: the maximal rate of RuBisCO carboxylation (V_cmax_) and the maximal rate of electron transport (J_max_)^18^. Consistent with a decrease in CO_2_ assimilation, values for both parameters were reduced significantly in *Svshr2* plants compared to wild type (Fig. 4f and g). Altogether, mutants had impaired photosynthetic capacity and a defective C4 cycle, likely as a consequence of a downregulated photosynthetic gene expression program and changes in bundle sheath cell function.

### Seed yield decreases under heat stress

Because plant biomass production and seed yield are frequently correlated with photosynthetic rates, we evaluated if mutant plants had decreased yield compared to wt. We grew all genotypes, except for *Svshr1/2* double mutants which never set any seeds, in two different conditions: temperate (23°C) with moderate light intensity (200 μmol photons m^−2^ s^−1^), versus high temperature (30°C) and saturating light (≈ 1100 μmol photons m^−2^ s^−1^). All genotypes experienced heat stress at 30°C because plants (including wild type) were visibly smaller and produced less leaves (Fig. 5a, b and c). However, heat stress clearly had a more severe effect on *Svshr2* since values for biomass and seed yield (measured as total grain weight per plant) significantly decreased compared to wild type (Fig. 5, b &d). Importantly, these effects cannot be attributed to root growth defects because *Svshr2* plants developed normal roots (Extended Data Fig.4). Moreover, when heat stress was removed there were no significant biomass or yield differences across genotypes (Fig. 5, c & e). We also quantified the number of seeds per panicle and found they were similar for all genotypes (Extended Data Fig. 5), suggesting that decrease in yield was probably due to incomplete filling.

**Fig. 5.**
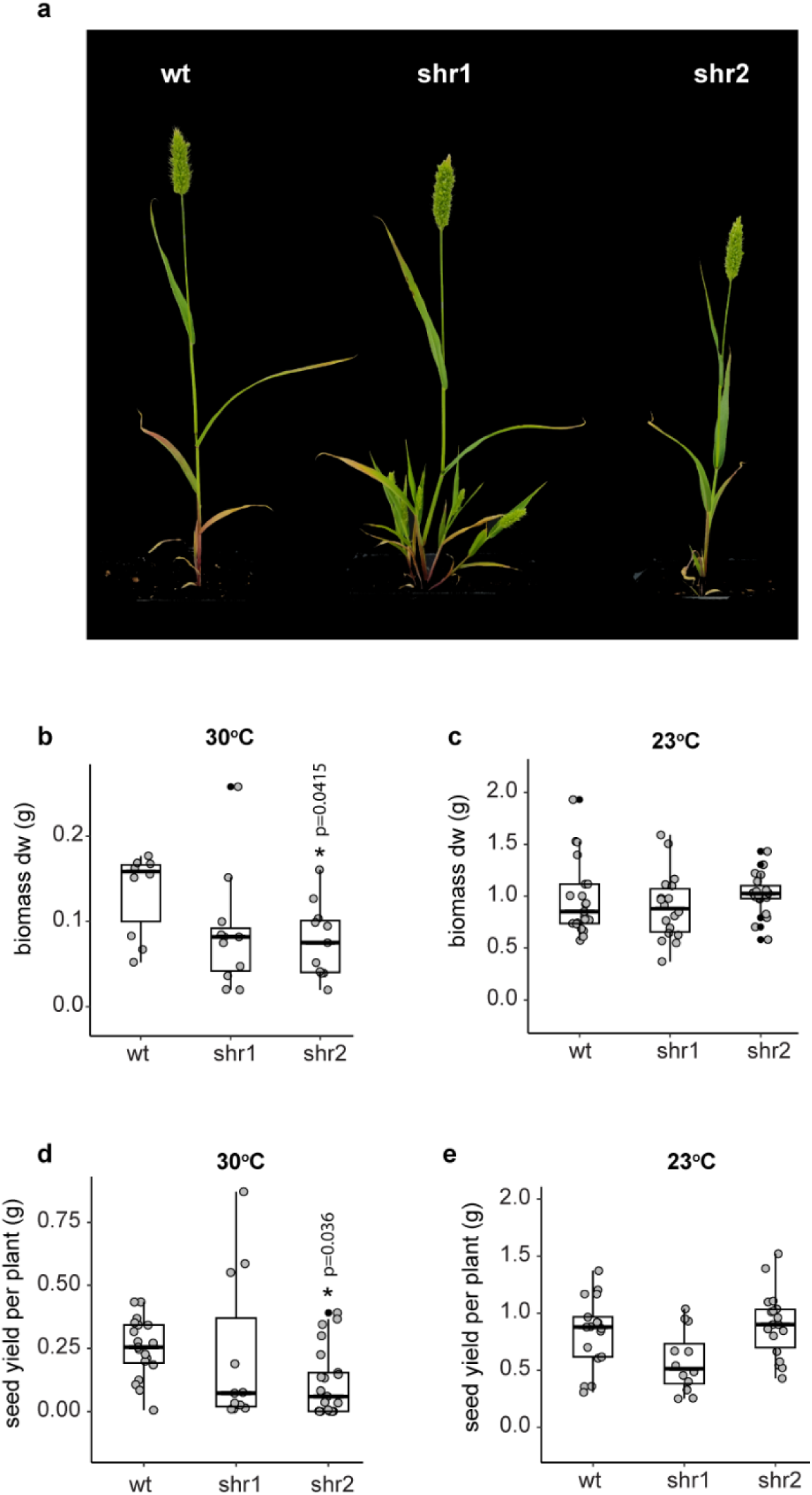
Quantification of plant biomass and yield. (**a**) Representative images of plants with fully developed inflorescences grown at 30 °C. (**b**) Above ground biomass of plants grown at 30°C measured as dry weight for wt (n = 10), Svshr1 (n = 11), and Svshr2 (n = 11). (**c**) Above ground biomass of plants grown at 23°C, measured as dry weight for wt (n = 21), Svshr1 (n = 18), and Svshr2 (n = 19). (**d**) Seed yield per plant grown at 30 °C for wt (n = 19), Svshr1 (n = 11), and Svshr2 (n = 21). (**e**) Seed yield per plant grown at 23 °C for wt (n = 17), Svshr1 (n = 12), and Svshr2 (n = 18). Asterisks indicate statistically significant difference with respect to wt (p < 0.05, Dunnet test after one way analysis of variance on all genotypes). Box plot center lines represent the median and box limits indicate the 25^th^ and 75^th^ percentiles.

The analysis of the effects of heat on physiology are highly consistent with SHR having a major role in Kranz anatomy and bundle sheath cell identity. These traits evolved to improve carbon assimilation under high temperatures and do not represent an advantage under cooler conditions^19^. Therefore, plants with a compromised C4 cycle are expected to show temperature-dependent phenotypes as observed in *Svshr2*. Taken together, this result supports a model in which *SHR* plays a major role in *Setaria’s* ability to carry out the C4 cycle.

Thus, by controlling key aspects of C4 anatomy, as well as promoting the unique C4 gene expression programs and enzymatic pathways involved in the exchange of four carbon chains to the bs, *SHR* ultimately contributes to modulate susceptibility to high heat – precisely the environmental condition under which C4 photosynthesis evolved.

## Discussion

We took an integrative approach combining histological, single-cell genomics and physiological analyses to study the role of *SHORT ROOT* in C4 photosynthesis. In agreement with previous studies^7,9^, we determined that *SHR* is involved in leaf patterning by regulating mesophyll cell divisions and vein density. However, we discovered that in *Setaria SHR* also has a crucial role in the development and placement of rank2 veins, which are specific to C4 species. We observed that wild type plants have a repetitive pattern in which each lateral vein is separated by up to six rank2 veins (Fig. 1c). This pattern changed substantially in both *Svshr2* single and *Svshr1/2* double mutant plants, decreasing rank2 vein density (Fig. 1k). Importantly, neither rank1 nor lateral vein development was negatively affected. In fact, their density increased (Fig. 1, j & l). A possible interpretation of this vein-type specific effect is that SHR may have a role in procambium initiation. Procambium gives rise to vascular centers in both C3 and C4 plants. However, in C4 plants the cells in the innermost layer remain competent to form procambium for longer, so that extra files of small veins can develop and fill the space between existing veins^4^. Hence, defects in maintaining procambium competence are expected to affect development of rank2 veins, which are the last to be specified, without affecting development of lateral veins that are produced first. Although this is in line with our observations, more histological and molecular analyses are needed at earlier developmental time points to confirm a possible role in procambium maintenance.

In addition to its role in vein patterning, *SHR* has been implicated in bundle sheath development in *Arabidopsis thaliana*. Evidence comes from leaf sections of mutant plants in which the bundle sheath layer appears to be missing^20^. However, this has not been confirmed at the molecular level. Moreover, because *Arabidopsis* is a C3 plant, it is unclear if *SHR* may have a similar function in C4 species. Our results indicate that there are differences in cell morphology suggesting cell fate specification defects. Through high resolution transcriptomics, we quantified cell identity changes and found that *SHR* is important for both maintaining bundle sheath identity and repressing mesophyll cell fate (Fig. 2f). Notably, it has been shown that in several C4 grasses bundle sheath and mesophyll cells originate from the same cell lineage, meaning that early in development they acquire distinct identities based on positional signals^21^. Based on our results, SHR may function as a fate specification factor that is needed for the differentiation of these cell types. It is important to note however, that gene expression programs associated to bundle sheath identity should differ considerably in C3 and C4 plants. Because this tissue acquired photosynthetic functions in the latter, C4 bundle sheath differentiation should involve the expression of photosynthetic genes.

In line with this, we found significant changes in the expression of many genes related to photosynthesis regulation, chlorophyll biosynthesis and chloroplast organization in mutant bundle sheath cells. This effect was also observed in photosynthetic mesophyll but not in other cell types (Fig. 2e). Importantly, downregulation of photosynthesis genes clearly impacted plant physiology via reduced photosynthetic rates in both *Svshr2* single and *Svshr1/2* double mutants (Fig. 4). Although *SHR* has never been directly associated with photosynthesis regulation, a recent study demonstrated that *SCR* double mutants in maize have less chlorophyll, as well as reduced maximum photosynthetic capacity (A_max_)^22^. Therefore, a logical hypothesis is that SHR could regulate the expression of photosynthetic genes through its putative interaction with *SCR*. However, *Setaria SHR* mutants show stronger photosynthesis deficient phenotypes: In contrast to SCR mutants, a phenotype is already evident in both *Svshr1* and *Svshr2* single mutants. Furthermore, double mutant plants have no detectable CO_2_ assimilation rates. Hence, *SHR* probably affects photosynthesis through *SCR* dependent and independent mechanisms. Supporting this hypothesis is the fact that *SHR* mutants have increased CO_2_ compensation points, which is not observed in *SCR* maize mutants (Fig. 4, c & d). This indicates that *SHR* is necessary for the normal operation of the C4 cycle.

We showed that defects at the molecular and anatomical level had important consequences for plant growth and productivity. Importantly, data shows that the most severe pleiotropic effects of perturbing SHR function do not account for key phenotypes related to C4 photosynthesis. Even though double mutants were severely stunted, plants with the *Svshr2* mutant allele grew normal size shoots and roots. Moreover, like most C4 traits, productivity defects in *Svshr2* were temperature dependent. Thus, we conclude that *SHR* regulates developmental and physiological traits in *Setaria* necessary for the emergence of C4 syndrome.

## Methods

### Accesion numbers Setaria

SHR

Sevir.9G361300 (SvSHR1)

Sevir2G383300 (SvSHR2)

### Plant growth conditions

For protoplast generation, Setaria seeds were incubated in 3% (v/v) sodium hypochlorite for 8 minutes and washed at least four times with sterile water. Seeds were germinated on square plates containing 0.5X Murashige and Skoog (MS) medium. Plates were kept in a Percival high light chamber for 24 days with a dark-light cycle of 16 h light and 8 h dark at 23 °C and 50% RH. For photosynthesis measurements, histological analyses, and root length quantification, Setaria plants were grown as above for 28-30 days. Plants used for seed yield quantification were grown at two different conditions in soil substrate: a greenhouse with an average temperature of 30oC, 13 hours light (1100 μmol photons m−2 s−1) and 11 hours dark, and in a walk-in plant chamber at 23oC with the same light/dark cycle, but at a lower light intensity (200 μmol photons m−2 s−1). They were grown for 8 weeks or until seeds were completely mature. Plants were not exposed to water stress. For half-mount in-situ hybridization and suberin staining, seeds were grown on solid substrate (vermiculite, perlite, and organic soil) under long-day conditions with a stable temperature of 27°C.

### Leaf histological sections

The third fully expanded leaf was removed from 28 day old plants (i.e. first basipetal leaf) and used for histological sections. Segments of the leaf encompassing the midrib were cut from the midpoint along the proximal/distal axis and fixed in 2 µL FAE solution (3.7% paraformaldehyde with 5% acetic acid glacial and 50% absolute ethanol). Vacuum (0.6 bar) was applied for 20 minutes, after which the leaf sections were rinsed and store at 4°C in 70% ethanol (v/v). Samples were dehydrated through 80%, 90%, and 96% (v/v) ethanol for 1 h each. Dehydrated leaf sections were then incubated in 1:1 SF10-Basic Solution Technovit 7100/ ethanol (v/v) for 2 h. After this incubation period, SF10-Basic Solution Technovit 7100 and Hardener I (infiltration solution) were applied and vacuumed for 40 min. Fixed tissue was then embedded in 15:1 infiltration solution/ Hardener II (polymerization solution I) and incubated for 48 hours. Finally, polymerization solution 2 (Technovit 3040 kit) was added. Once cooled, blocks were trimmed, and sections (10 μm) were cut using a microtome and placed on slides to dry overnight. Slides were stained in 0.05% (w/v) Toluidine Blue (50 mM citrate buffer, pH 4.4) for 5 s, rinsed in distilled H2O, and then dried and covered with a coverslip. Leaf segments were imaged using a Leica LSM 800 confocal microscope. Images were taken using bright-field and fluorescence illumination. Cell wall autofluorescence was captured using 408 nm laser, and collection at 495-515 nm.

### Protoplast generation and single-cell RNAseq

Leaf segments were sectioned in ∼3-mm fragments using a scalpel. Fragments were placed in a small petri dish and weighted to calculate protoplast yield. Protoplasting solution was added immediately after weighting, which consisted of 1.25% (w/v) Cellulase R-10, 2% (w/v) Cellulase RS, 0.4% (w/v) macerozyme R10, and 0.2% (w/v) pectolyase Y23. Tissue digestion was carried out with gentle shaking at room temperature for 1 hour and 40 minutes. The cell suspension was then filtered using a 40 μm nylon filter and cells were then centrifuged at 200 rcf for 5 min. Cells were resuspended in pre-filtered (0.22 µm) washing buffer (0.4 M mannitol, 0.02 M MES, 0.02 M KCl, 0.01 M CaCl2, 0.015 mM BSA). A cell aliquot was stained with fluorescein diacetate (2µg/µl) for 3 min and counted in a hemocytometer to determine cell concentration and viability. Cells were then re-suspended at a concentration of 2000-3000 cells/μL and loaded into a Single Cell A chip (10× Genomics). Approximately 10,000 cells were loaded per replicate. Three independent replicates per genotype were performed. Chips were loaded on a Chromium Controller (10x Genomics) to generate single-cell GEMs. Single-cell libraries were then prepared using the Chromium Single Cell 3’ library kit, following manufacturer instructions. The quality of resulting DNA libraries was assessed with an Agilent TapeStation system. Library concentration was determined by quantitative PCR (qPCR) and sequenced with an Illumina Novaseq platform using a 1x150 configuration. Raw scRNAseq data was analyzed by Cell Ranger 5.0.1 (10x Genomics) to generate gene-cell matrices. Gene reads were aligned to Setaria viridis 2.1 reference genome.

### Data integration and UMAP analysis

Cell-type analysis and clustering were performed using Seurat v4.3.0^1^. To remove low quality cells we applied three filters: first, genes with counts in fewer than three cells were excluded from the analysis and their counts were removed. Second, cells with fewer than 250 genes were removed. Third, cells with more than 5% counts mapping to mitochondrial genes were excluded. Next, all replicates from wt and shr2 mutant were integrated for downstream analyses as follows: counts were log-normalized with the LogNormalize function and a subset of genes that showed large cell-to-cell variation in the data set was calculated by directly modeling the mean-variance relationship with the VST function. We used the FindIntegrationAnchors function to set anchors across replicates and genotypes, using 20 dimensions. An integrated expression matrix containing cells from wt and shr2 replicates was produced with the IntegrateData function. Dimensionality reduction of the integrated expression matrix was performed by scaling the data (linear transformed) using the ScaleData function and calculating Principal Components (PCA). 15-20 main components were chosen. Cells were then clustered using the FindNeighbor function to create a K nearest neighbor (KNN) graph followed by the FindClusters function with a resolution of 0.6. These clusters were used as input for nonlinear dimensional reduction with the Uniform Manifold Approximation and Projection (UMAP) method using 20 principal components.

### Cell annotation and differential expression analyses

To determine cluster identity, we searched for leaf cell-type specific marker genes supported by either experimental or single-cell expression data in the PlantCellMarker^2^ portal (https://www.tobaccodb.org/pcmdb/homePage). Since there is no information on Setaria, we used markers identified in maize and looked for Setaria orthologs using the Monocots Plaza 5.0 integrative method. We further confirmed the identity of bundle sheath and mesophyll clusters by fluorescent in-situ hybridization of cluster marker genes identified from our dataset using the ROC algorithm (Seurat).

For differential expression analyses between wt and shr2, we performed a psedobulk analysis in which counts from all cells with the same identity (cell type) and replicate were aggregated with the AggregateExpression function in Seurat. The pseudobulked expression matrix was then used to calculate DEGs using the DESeq2 package^3^. Gene Ontology analyses were then performed on wt and shr2 cell type DEGs using the Monocots Plaza 5.0 GO analysis with the Bonferroni test.

### ICI calculation

The Index of Cell Identity (ICI) was calculated for each cell as described before^4^. In brief, DEGs for each tissue were identified using DESeq2 with wt pseudobulked cell cluster counts as input. Next, specificity values were calculated on these genes using the Spec algorithm^5^. This algorithm ponders the global expression of each gene across all cell types, as well as the effect of noise, to identify genes with high specificity for a given cell type. Calculation of optimal bin size for each gene was empirically attributed and performed according to Effroni et al. (2016). The top 100 markers (based on wt Spec scores) were selected for each tissue and used to calculate ICI scores for both wt and shr2 cells. Scores were visualized as density plots.

### Half-mount in situ hybridization

Recently emerged leaves from 4-week-old plants were collected and placed in FAA fixative solution (4% formaldehyde, 5% glacial acetic acid, 50% ethanol in DEPC water) for 2 hours under gentle vacuum (-0.5 kPa). Excess FAA was removed, and hand sectioning across the longitudinal axis of the leaves was performed with a scalpel blade under a stereoscope. The cross-sectioned samples were placed again in FAA solution overnight at 4°C. Next, samples were treated with a series of washes in ethanol at increasing concentrations (70%, 90%, 100%) at room temperature (25°C) before being stored in 100% methanol at -20°C. HCR RNA-FISH hybridization and amplification were performed according to a recent half-mount protocol tested in monocots^6^. Before sample visualization, samples were incubated in Calcofluor White 0.1% (w/v) for 1 min to stain the cell wall. Samples were gently inserted between agarose gel plates using fine forceps and transferred onto a glass slide, covered with a coverslip, and imaged on a Leica LSM800™ confocal microscope. Probes were detected using 561 nm excitation and 565-578 nm collection wavelengths. Cell membrane autofluorescence was detected using 405 nm excitation and 410-450 nm wavelengths. The brightness and contrast were adjusted by FIJI software. Probes with split initiator B1 for the target genes were designed and manufactured by Molecular Instruments. HCR reagents, including hairpins with a B1 amplifier sequence and the fluorescent dye Alexa Fluor 546 were purchased from Molecular Instruments.

### Suberin staining

S. viridis leaves were collected three weeks after germination and dehydrated in a series of washes with ethanol at increasing concentrations (70%, 90%, 100%) at room temperature (25°C) before being stored in 100% methanol at -20°C. Next, leaf cross sections were generated with a scalpel under a stereoscope. For suberin staining, the cross-sectioned samples were incubated in a freshly prepared solution of 0.01% (w/v) Fluorol Yellow 088 (Santa Cruz Biotechnologies) in lactic acid at 70°C for 1 hour2. The samples were washed two to three times with water and then incubated in Calcofluor White 0.1% (w/v) for 1 minute to stain the cell wall. Samples were gently inserted between agarose gel plates using fine forceps and transferred onto a glass slide, covered with a coverslip, and imaged on a Leica LSM800™ confocal microscope. Fluorol Yellow signal was detected using 488 nm excitation and 540-570 nm collection wavelengths. Cell membrane autofluorescence was detected using 405 nm excitation and 410-450 nm wavelengths. The brightness and contrast were adjusted by FIJI software.

### Photosynthesis measurements

Steady state Amax measurements and A/Ci curves were generated using an LI-6800 (LI-COR) photosynthesis measuring system, containing a custom-made LED white light lamp with a color temperature similar to sunlight^7^. Amax measurements were made on fully expanded leaves at 800 μmol/mol CO2, 1800 μmol photons m−2 s−1, 60% humidity and 26°C leaf temperature. Leaves were allowed to acclimate until photosynthetic rate had stabilized for at least 30 min at 400 μmol/mol CO2. A/Ci curves were generated by measurement of photosynthetic rates at 400 (omitted from curve plotting), 1000, 800, 600, 400, 300, 200, 100, 50, 25, and 10 μmol/mol CO2, at this point Amax was recorded. Compensation points were estimated by fitting a polynomial trendline of the modelled data and solving this to estimate the x-axis intercept and thus the CO2 compensation point.

### Biomass measurements

Upon reaching maturity, the plant panicles were removed and the aerial part of the plant was collected at the soil level. Subsequently, the harvested samples were carefully covered in aluminum sheets and subjected to desiccation in an oven at 65°C for a minimum period of 72 hours. The individual dried samples were then weighted using an electronic scale (Pioneer PX84). Dunnett’s statistical test after one-way anova was employed to assess significant differences between genotypes and treatments.

### Seed harvest and quantification

5-week-old plants were monitored to assess initiation of panicle production. After panicle emergence, each panicle was enclosed with a protective organza bag, which remained in place until seed harvest to prevent seed loss. Upon complete senescence of the plants, the inflorescence was severed at its base. Subsequently, all seeds adhering to the panicle were carefully extracted by gently compressing the dried inflorescence. Pictures of seeds from individual panicles were taken and used to determine seed count with the ImageJ software. The cell counter function was used to distinguish between seeds and inflorescence debris. Seed weight was determined by placing all seeds from individual panicles in analytic balance (Pioneer PX84). The Dunnett test after one-way anova was employed to evaluate differences in seed production. Statistical analysis was carried out using RStudio.

### Root length Measurements

4-week-old plants were carefully removed from the agar medium taking care not to damage or severe the roots. They were placed on wet germination paper and used to take pictures for further analysis. Roots were measured using the ImageJ software. Dunnett’s statistical test was applied to evaluate significant differences between treatments. The Dunnett test after one-way anova was employed to evaluate differences in length. Statistical analysis was carried out using RStudio.

### Statistical analysis

To compare wt versus mutant samples we applied a one-way ANOVA analysis of variance followed by Dunnett’s test. In all cases the means of mutant genotypes were compared to that of wild type samples, which was used as a control group. This test allows to control for type I error rate. Calculation was performed using programing language R with 95% confidence intervals and a *P* value equal or less than 0.05. Polynomial regression fits and box plots were also calculated with the same software. All measurements used to perform statistical tests were taken from distinct samples.

## Acknowledgements

**Funding:** C.O.R. and D.C.E. were supported by SEP-CONACYT grant I0017. C.A.G., E.P.F., J.C.R., are supported by CONHACYT postgraduate studies grant. M.H.C. is supported by CZI grant 2021-240108. K.D.B. and B.G. are supported by the National Institutes of Health grant R35GM136362. **Author contributions**: D.C.E. performed all histological, physiological and plant yield experiments. D.C.E. and C.O.R. designed the experiments. C.O.R. wrote the manuscript. S.F., K.D.B., M.H.C., C.O.R., conceived the project and guided the experiments. C.A.G. performed single-cell RNAseq experiments. E.P.F. Carried out in situ hybridization experiments and image analyses. B.G. and M.H.C assisted in the generation and analysis of single-cell RNA data. A.V.R. assisted in the generation of photosynthesis measurements. J.C.R. and V.C.B. assisted in histological image analyses and measurements.

## Supplementary information

**Extended Data Fig. 1.**
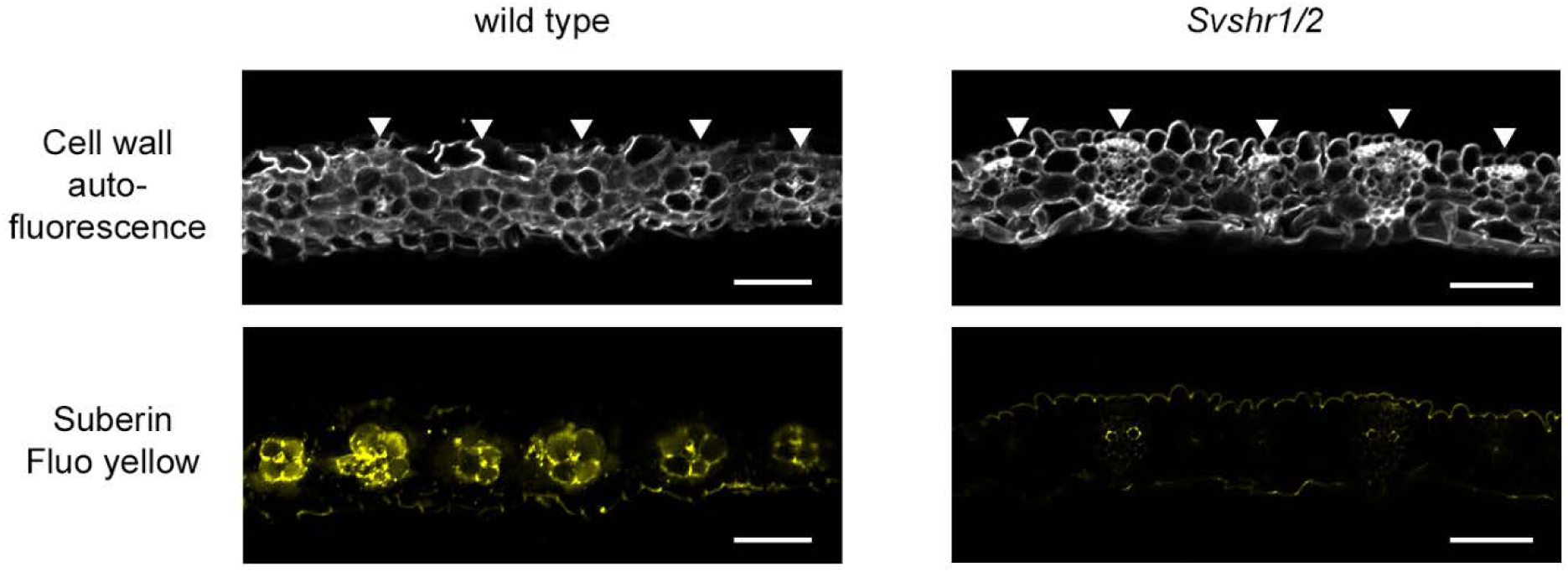
Presence of suberin in bundle sheath cells from wt and *shr1/2*. Suberin was stained with the fluorescent dye fluorol yellow 088. Signal is clearly detected in the cell wall of all bundle sheath cells and some vascular cells in wt samples (left panel). In contrast, no signal was detected in the bundle sheath cell wall of shr1/2 mutant samples (right panel). Arrowheads point to the position of vascular bundles. Yellow coloration corresponds to fluorol yellow signal. Grey corresponds to cell wall autofluorescence. Scale bars, 50 µm.

**Extended Data Fig. 2.**
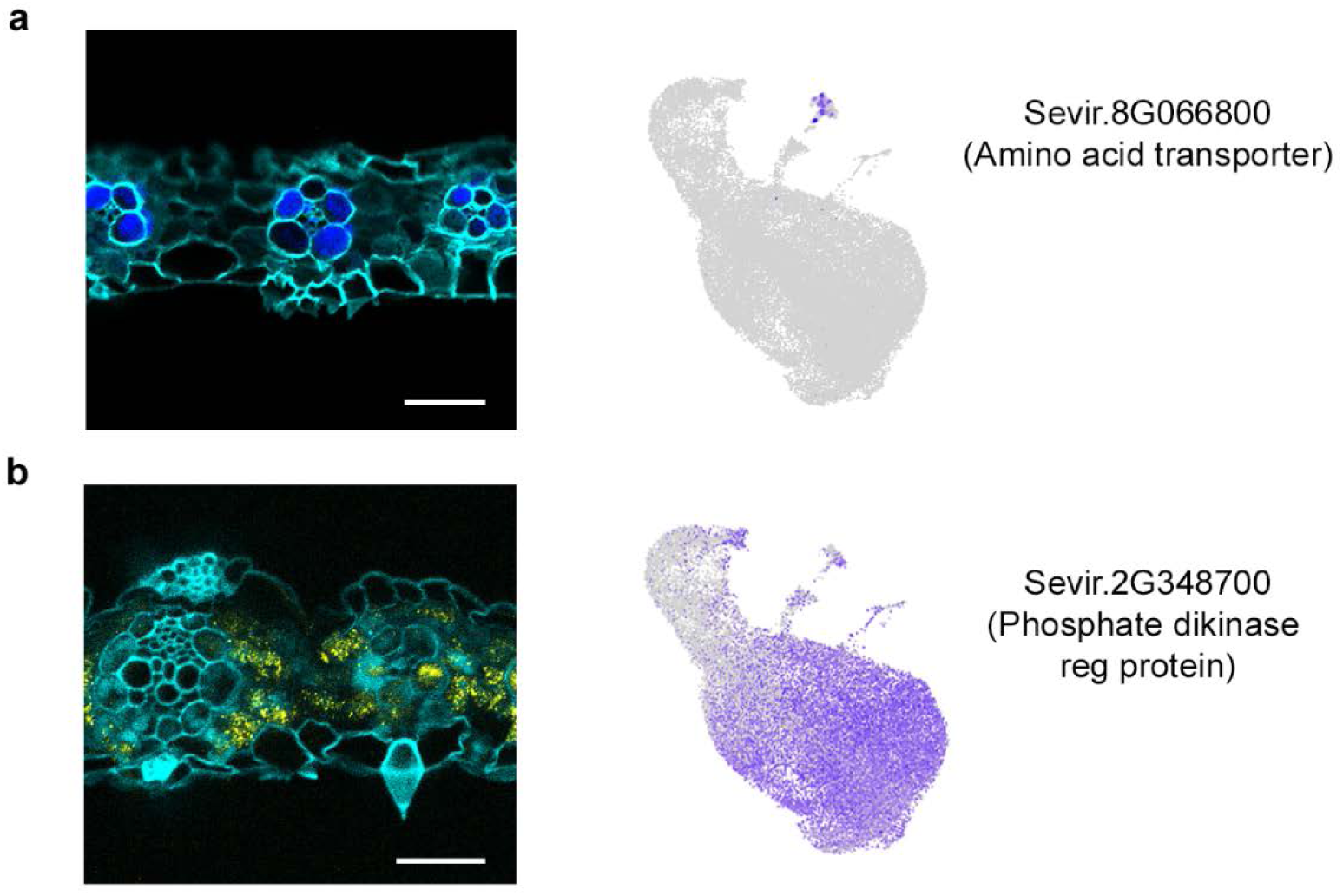
Transcript localization of bundle sheath and mesophyll marker genes. (**a**) *In situ* hybridization (RNA FISH) showing the localization of Sevir.8G066800, an aminoacid transporter found to be uniquely expressed in the bundle sheath cluster (right). (**b**) In situ hybridization showing the localization of Sevir.2G348700, a phosphate dikinase regulatory protein found to be mostly expressed in mesophyll clusters (right). Blue and yellow correspond to bundle sheath and mesophyll marker signal, while cyan signal corresponds to cell wall autofluorescence. Scale bars, 25 µm.

**Extended Data Fig. 3.**
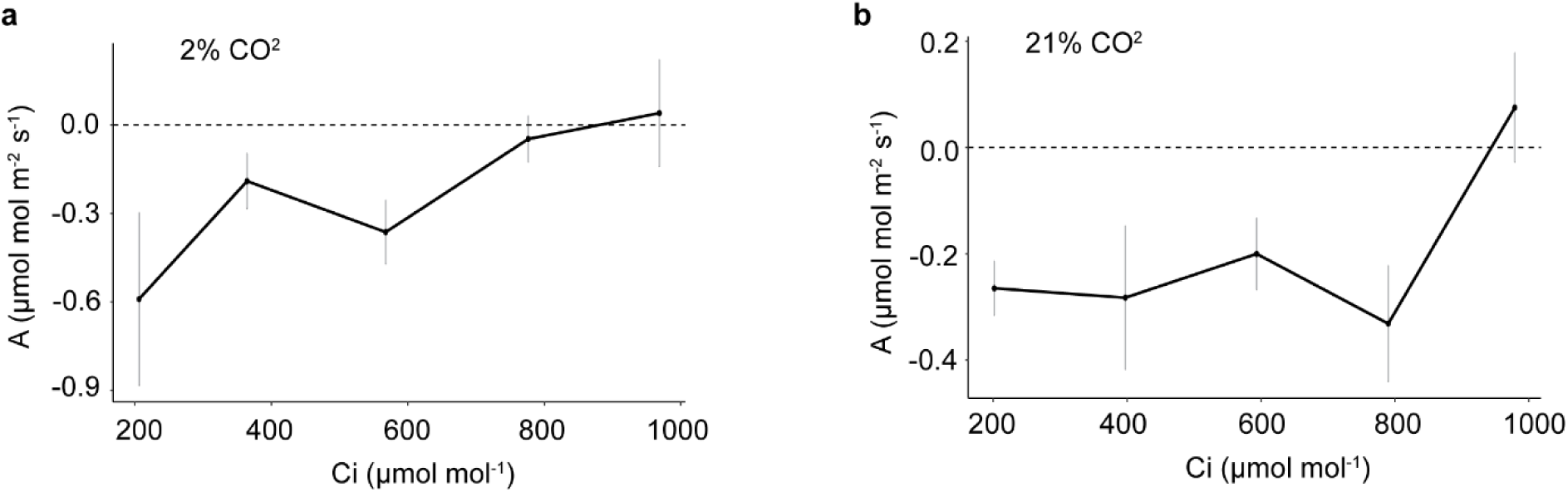
Photosynthetic rates in shr1/2 double mutant plants. (**a**) A/Ci curves of *shr1/2* mutants (*n* = 6) at 2% oxygen. (**b**) A/Ci curves of *shr1/2* mutants (*n* = 6) at 21% oxygen. Error bars (grey) are standard deviation of the mean.

**Extended Data Fig. 4.**
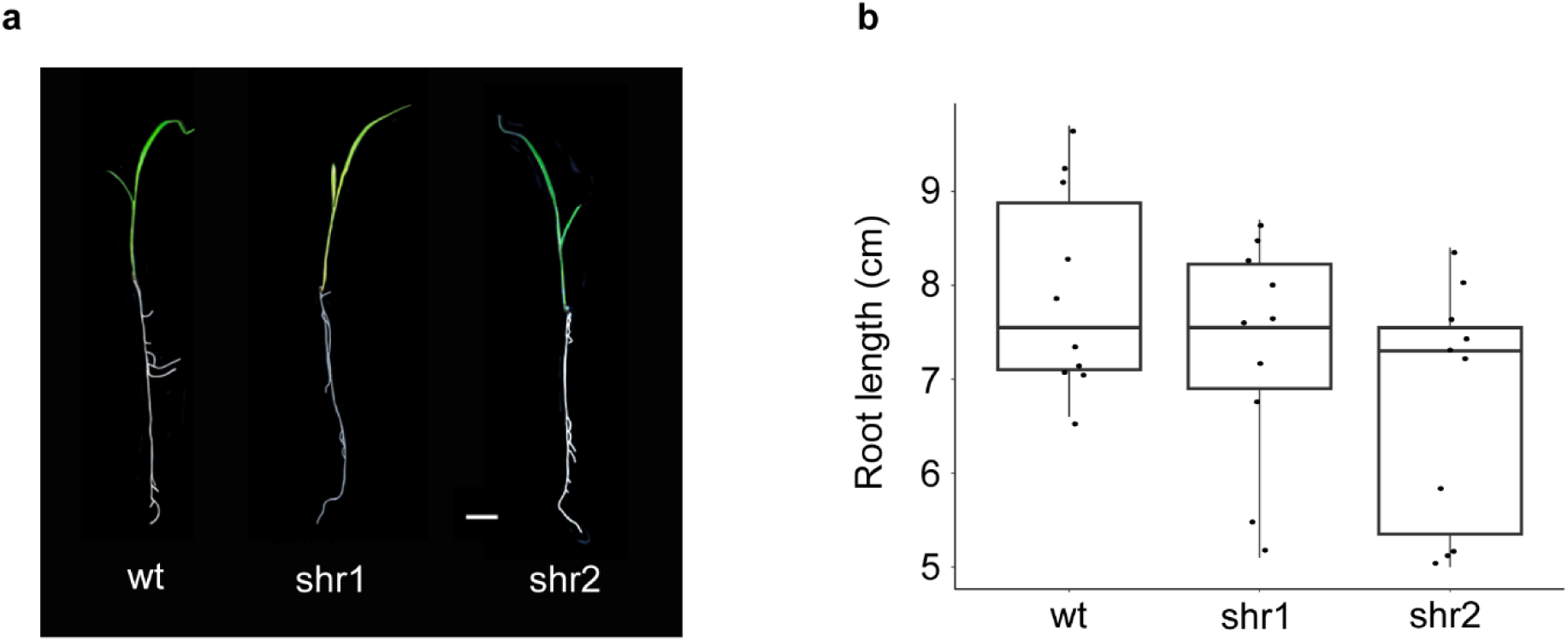
Root length in mutant genotypes. (**a**) Representative image of 3-week-old plants showing root length and morphology. (**b**) Measurements of root length from wt (n=10), Sv*shr1* (n = 10), and *Svshr2* (n=10) plants. No significant differences across genotypes were found after a one-way analysis of variance followed by Dunnet test on mutant and wt genotypes. Box plot center lines represent the median and box limits indicate the 25^th^ and 75^th^ percentiles.

**Extended Data Fig. 5.**
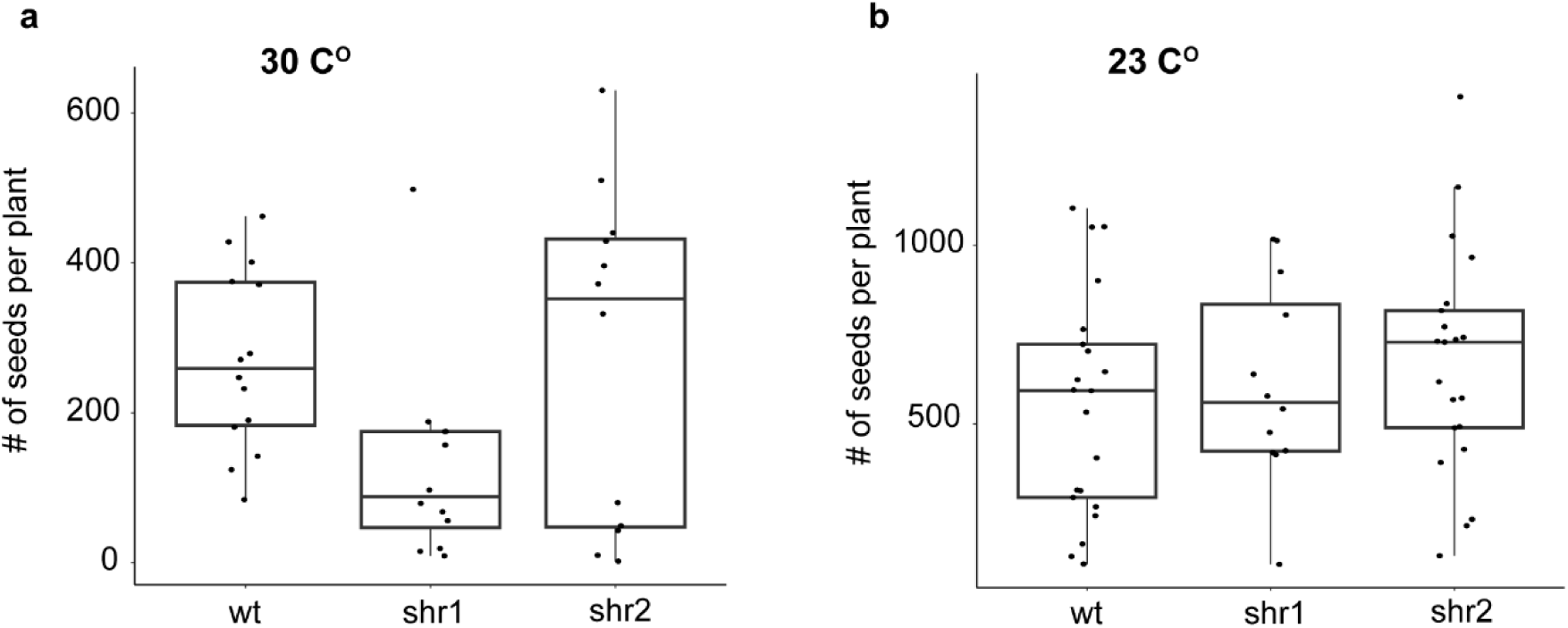
Number of seeds per plant. (**a**) Quantification of the number of seeds per plant from wt (n=14), *Svshr1* (n=11), and *Svshr2* (n=12) genotypes grown at 30 C° and saturated light conditions. (**b**) Quantification of the number of seeds per plant from wt (n=21), *Svshr1* (n=12), and *Svshr2* (n=22) genotypes grown at 23 C° and low light. No significant differences across genotypes were found after a one-way analysis of variance followed by Dunnet test on mutant and wt genotypes. Box plot center lines represent the median and box limits indicate the 25^th^ and 75^th^ percentiles.

**Supplementary Table 1.**
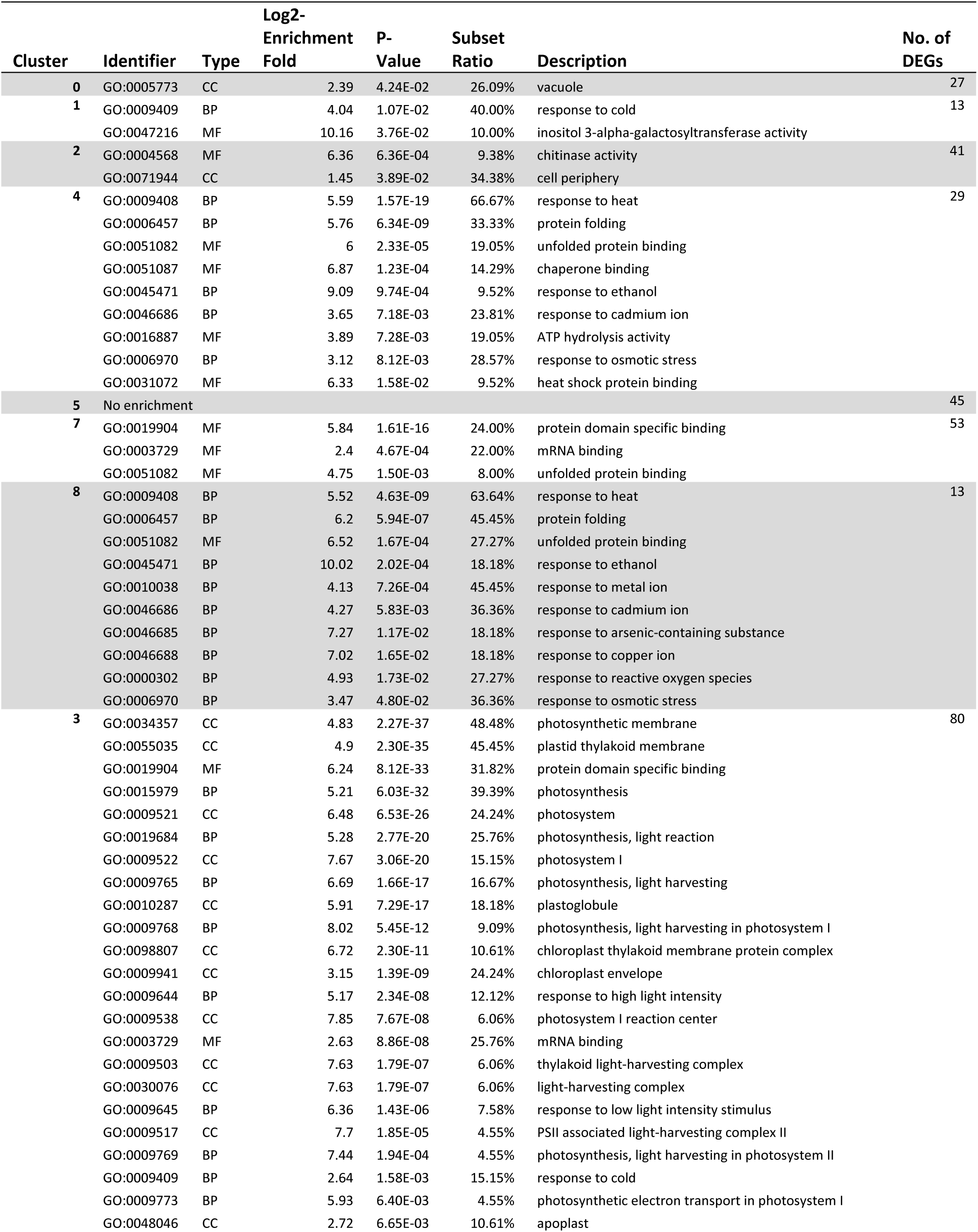
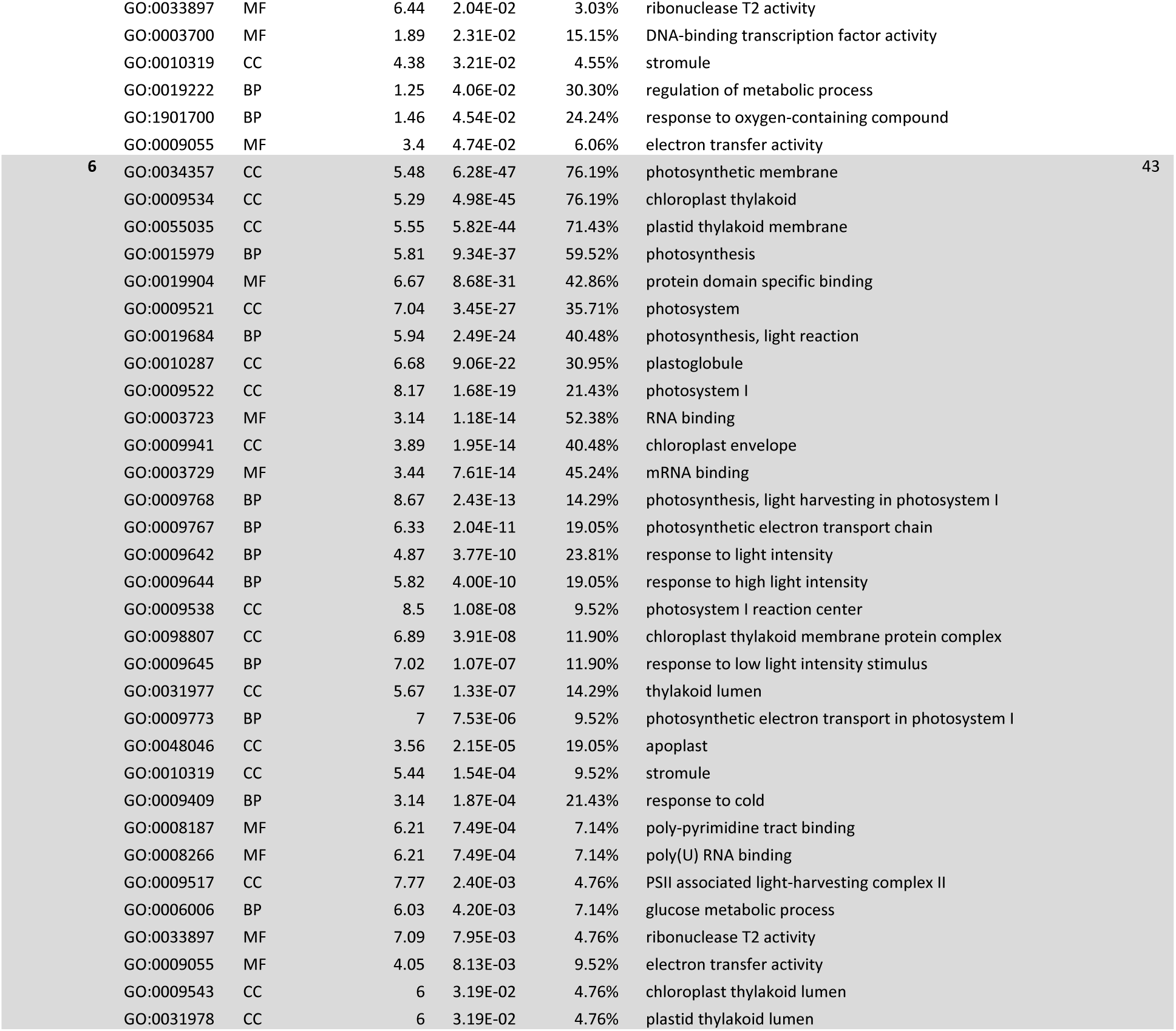
Gene ontology enrichment of individual mesophyll cluster DEGs. Clusters enriched in photosynthetic functions are grouped at the bottom.

| Cluster | Identifier | Type | Log2-Enrichment Fold | P-Value | Subset Ratio | Description | No. of DEGs |
| --- | --- | --- | --- | --- | --- | --- | --- |
| 0 | GO:0005773 | CC | 2.39 | 4.24E-02 | 26.09% | vacuole | 27 |
| 1 | GO:0009409 | BP | 4.04 | 1.07E-02 | 40.00% | response to cold | 13 |
|  | GO:0047216 | MF | 10.16 | 3.76E-02 | 10.00% | inositol 3-alpha-galactosyltransferase activity |  |
| 2 | GO:0004568 | MF | 6.36 | 6.36E-04 | 9.38% | chitinase activity | 41 |
|  | GO:0071944 | CC | 1.45 | 3.89E-02 | 34.38% | cell periphery |  |
| 4 | GO:0009408 | BP | 5.59 | 1.57E-19 | 66.67% | response to heat | 29 |
|  | GO:0006457 | BP | 5.76 | 6.34E-09 | 33.33% | protein folding |  |
|  | GO:0051082 | MF | 6 | 2.33E-05 | 19.05% | unfolded protein binding |  |
|  | GO:0051087 | MF | 6.87 | 1.23E-04 | 14.29% | chaperone binding |  |
|  | GO:0045471 | BP | 9.09 | 9.74E-04 | 9.52% | response to ethanol |  |
|  | GO:0046686 | BP | 3.65 | 7.18E-03 | 23.81% | response to cadmium ion |  |
|  | GO:0016887 | MF | 3.89 | 7.28E-03 | 19.05% | ATP hydrolysis activity |  |
|  | GO:0006970 | BP | 3.12 | 8.12E-03 | 28.57% | response to osmotic stress |  |
|  | GO:0031072 | MF | 6.33 | 1.58E-02 | 9.52% | heat shock protein binding |  |
| 5 | No enrichment |  |  |  |  |  | 45 |
| 7 | GO:0019904 | MF | 5.84 | 1.61E-16 | 24.00% | protein domain specific binding | 53 |
|  | GO:0003729 | MF | 2.4 | 4.67E-04 | 22.00% | mRNA binding |  |
|  | GO:0051082 | MF | 4.75 | 1.50E-03 | 8.00% | unfolded protein binding |  |
| 8 | GO:0009408 | BP | 5.52 | 4.63E-09 | 63.64% | response to heat | 13 |
|  | GO:0006457 | BP | 6.2 | 5.94E-07 | 45.45% | protein folding |  |
|  | GO:0051082 | MF | 6.52 | 1.67E-04 | 27.27% | unfolded protein binding |  |
|  | GO:0045471 | BP | 10.02 | 2.02E-04 | 18.18% | response to ethanol |  |
|  | GO:0010038 | BP | 4.13 | 7.26E-04 | 45.45% | response to metal ion |  |
|  | GO:0046686 | BP | 4.27 | 5.83E-03 | 36.36% | response to cadmium ion |  |
|  | GO:0046685 | BP | 7.27 | 1.17E-02 | 18.18% | response to arsenic-containing substance |  |
|  | GO:0046688 | BP | 7.02 | 1.65E-02 | 18.18% | response to copper ion |  |
|  | GO:0000302 | BP | 4.93 | 1.73E-02 | 27.27% | response to reactive oxygen species |  |
|  | GO:0006970 | BP | 3.47 | 4.80E-02 | 36.36% | response to osmotic stress |  |
| 3 | GO:0034357 | CC | 4.83 | 2.27E-37 | 48.48% | photosynthetic membrane | 80 |
|  | GO:0055035 | CC | 4.9 | 2.30E-35 | 45.45% | plastid thylakoid membrane |  |
|  | GO:0019904 | MF | 6.24 | 8.12E-33 | 31.82% | protein domain specific binding |  |
|  | GO:0015979 | BP | 5.21 | 6.03E-32 | 39.39% | photosynthesis |  |
|  | GO:0009521 | CC | 6.48 | 6.53E-26 | 24.24% | photosystem |  |
|  | GO:0019684 | BP | 5.28 | 2.77E-20 | 25.76% | photosynthesis, light reaction |  |
|  | GO:0009522 | CC | 7.67 | 3.06E-20 | 15.15% | photosystem I |  |
|  | GO:0009765 | BP | 6.69 | 1.66E-17 | 16.67% | photosynthesis, light harvesting |  |
|  | GO:0010287 | CC | 5.91 | 7.29E-17 | 18.18% | plastoglobule |  |
|  | GO:0009768 | BP | 8.02 | 5.45E-12 | 9.09% | photosynthesis, light harvesting in photosystem I |  |
|  | GO:0098807 | CC | 6.72 | 2.30E-11 | 10.61% | chloroplast thylakoid membrane protein complex |  |
|  | GO:0009941 | CC | 3.15 | 1.39E-09 | 24.24% | chloroplast envelope |  |
|  | GO:0009644 | BP | 5.17 | 2.34E-08 | 12.12% | response to high light intensity |  |
|  | GO:0009538 | CC | 7.85 | 7.67E-08 | 6.06% | photosystem I reaction center |  |
|  | GO:0003729 | MF | 2.63 | 8.86E-08 | 25.76% | mRNA binding |  |
|  | GO:0009503 | CC | 7.63 | 1.79E-07 | 6.06% | thylakoid light-harvesting complex |  |
|  | GO:0030076 | CC | 7.63 | 1.79E-07 | 6.06% | light-harvesting complex |  |
|  | GO:0009645 | BP | 6.36 | 1.43E-06 | 7.58% | response to low light intensity stimulus |  |
|  | GO:0009517 | CC | 7.7 | 1.85E-05 | 4.55% | PSII associated light-harvesting complex II |  |
|  | GO:0009769 | BP | 7.44 | 1.94E-04 | 4.55% | photosynthesis, light harvesting in photosystem II |  |
|  | GO:0009409 | BP | 2.64 | 1.58E-03 | 15.15% | response to cold |  |
|  | GO:0009773 | BP | 5.93 | 6.40E-03 | 4.55% | photosynthetic electron transport in photosystem I |  |
|  | GO:0048046 | CC | 2.72 | 6.65E-03 | 10.61% | apoplast |  |

|  |  |  |  |  |  |  |
| --- | --- | --- | --- | --- | --- | --- |
|  | GO:0033897 | MF | 6.44 | 2.04E-02 | 3.03% | ribonuclease T2 activity |
|  | GO:0003700 | MF | 1.89 | 2.31E-02 | 15.15% | DNA-binding transcription factor activity |
|  | GO:0010319 | CC | 4.38 | 3.21E-02 | 4.55% | stromule |
|  | GO:0019222 | BP | 1.25 | 4.06E-02 | 30.30% | regulation of metabolic process |
|  | GO:1901700 | BP | 1.46 | 4.54E-02 | 24.24% | response to oxygen-containing compound |
|  | GO:0009055 | MF | 3.4 | 4.74E-02 | 6.06% | electron transfer activity |
| 6 | GO:0034357 | CC | 5.48 | 6.28E-47 | 76.19% | photosynthetic membrane |
|  | GO:0009534 | CC | 5.29 | 4.98E-45 | 76.19% | chloroplast thylakoid |
|  | GO:0055035 | CC | 5.55 | 5.82E-44 | 71.43% | plastid thylakoid membrane |
|  | GO:0015979 | BP | 5.81 | 9.34E-37 | 59.52% | photosynthesis |
|  | GO:0019904 | MF | 6.67 | 8.68E-31 | 42.86% | protein domain specific binding |
|  | GO:0009521 | CC | 7.04 | 3.45E-27 | 35.71% | photosystem |
|  | GO:0019684 | BP | 5.94 | 2.49E-24 | 40.48% | photosynthesis, light reaction |
|  | GO:0010287 | CC | 6.68 | 9.06E-22 | 30.95% | plastoglobule |
|  | GO:0009522 | CC | 8.17 | 1.68E-19 | 21.43% | photosystem I |
|  | GO:0003723 | MF | 3.14 | 1.18E-14 | 52.38% | RNA binding |
|  | GO:0009941 | CC | 3.89 | 1.95E-14 | 40.48% | chloroplast envelope |
|  | GO:0003729 | MF | 3.44 | 7.61E-14 | 45.24% | mRNA binding |
|  | GO:0009768 | BP | 8.67 | 2.43E-13 | 14.29% | photosynthesis, light harvesting in photosystem I |
|  | GO:0009767 | BP | 6.33 | 2.04E-11 | 19.05% | photosynthetic electron transport chain |
|  | GO:0009642 | BP | 4.87 | 3.77E-10 | 23.81% | response to light intensity |
|  | GO:0009644 | BP | 5.82 | 4.00E-10 | 19.05% | response to high light intensity |
|  | GO:0009538 | CC | 8.5 | 1.08E-08 | 9.52% | photosystem I reaction center |
|  | GO:0098807 | CC | 6.89 | 3.91E-08 | 11.90% | chloroplast thylakoid membrane protein complex |
|  | GO:0009645 | BP | 7.02 | 1.07E-07 | 11.90% | response to low light intensity stimulus |
|  | GO:0031977 | CC | 5.67 | 1.33E-07 | 14.29% | thylakoid lumen |
|  | GO:0009773 | BP | 7 | 7.53E-06 | 9.52% | photosynthetic electron transport in photosystem I |
|  | GO:0048046 | CC | 3.56 | 2.15E-05 | 19.05% | apoplast |
|  | GO:0010319 | CC | 5.44 | 1.54E-04 | 9.52% | stromule |
|  | GO:0009409 | BP | 3.14 | 1.87E-04 | 21.43% | response to cold |
|  | GO:0008187 | MF | 6.21 | 7.49E-04 | 7.14% | poly-pyrimidine tract binding |
|  | GO:0008266 | MF | 6.21 | 7.49E-04 | 7.14% | poly(U) RNA binding |
|  | GO:0009517 | CC | 7.77 | 2.40E-03 | 4.76% | PSII associated light-harvesting complex II |
|  | GO:0006006 | BP | 6.03 | 4.20E-03 | 7.14% | glucose metabolic process |
|  | GO:0033897 | MF | 7.09 | 7.95E-03 | 4.76% | ribonuclease T2 activity |
|  | GO:0009055 | MF | 4.05 | 8.13E-03 | 9.52% | electron transfer activity |
|  | GO:0009543 | CC | 6 | 3.19E-02 | 4.76% | chloroplast thylakoid lumen |
|  | GO:0031978 | CC | 6 | 3.19E-02 | 4.76% | plastid thylakoid lumen |

**Supplementary Table 2.**
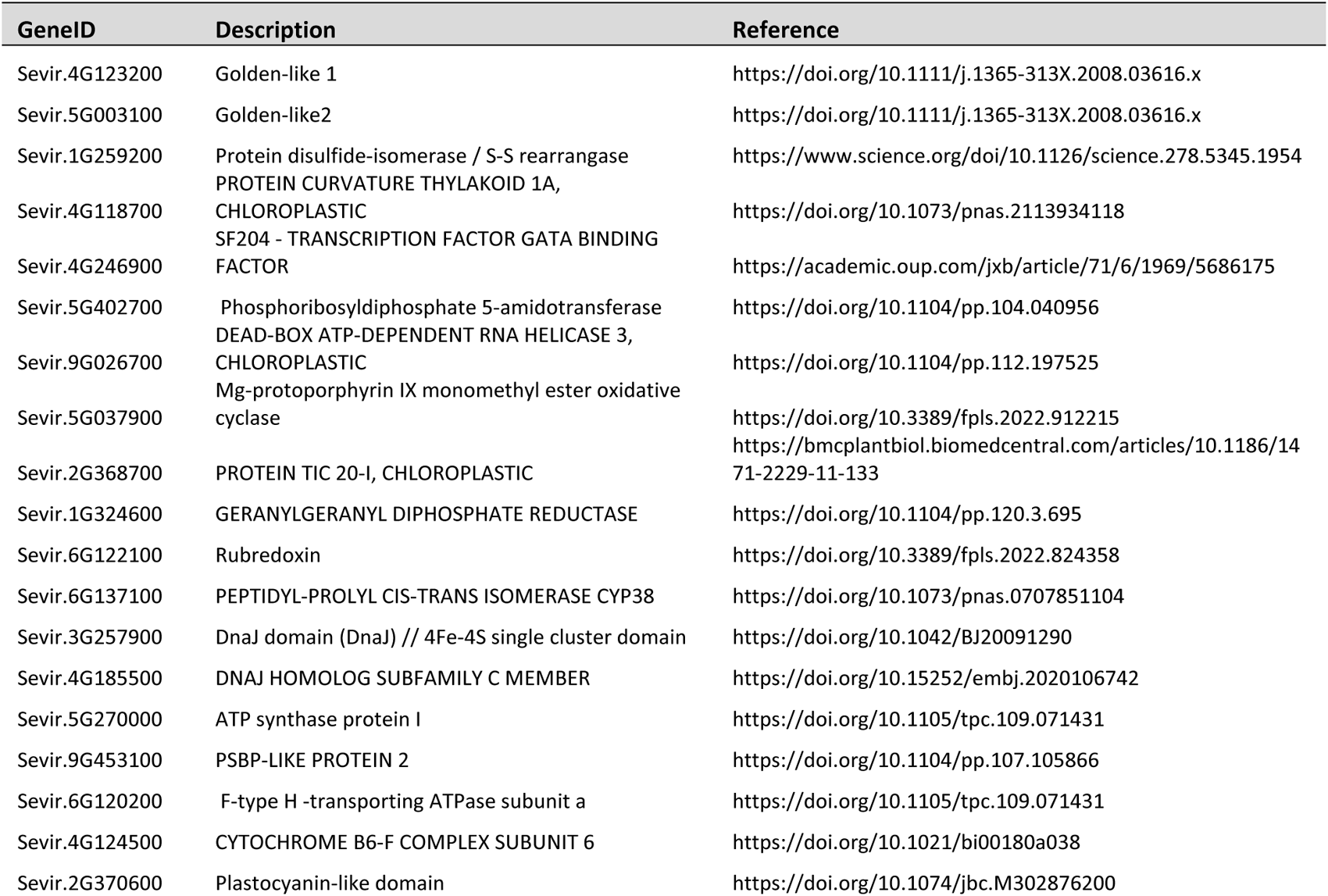
Downregulated chloroplast biogenesis and assembly genes in *shr2* bundle sheath.

## Data availability

Data has been deposited in the Gene Expression Omnibus (GEO) under accession GSE249836.

## References

1 Sage, R. F., Sage, T. L. & Kocacinar, F. Photorespiration and the evolution of C4 photosynthesis. Annu Rev Plant Biol 63, 19–47 (2012). 10.1146/annurev-arplant-042811-105511

2 Danila, F. R. et al. Bundle sheath suberisation is required for C(4) photosynthesis in a Setaria viridis mutant. Commun Biol 4, 254 (2021). 10.1038/s42003-021-01772-4

3 Griffiths, H., Weller, G., Toy, L. F. & Dennis, R. J. You’re so vein: bundle sheath physiology, phylogeny and evolution in C3 and C4 plants. Plant, cell & environment 36, 249–261 (2013). 10.1111/j.1365-3040.2012.02585.x

4 Sedelnikova, O. V., Hughes, T. E. & Langdale, J. A. Understanding the Genetic Basis of C4 Kranz Anatomy with a View to Engineering C3 Crops. Annual review of genetics 52, 249–270 (2018). 10.1146/annurev-genet-120417-031217

5 Edwards, G. E., Franceschi, V. R. & Voznesenskaya, E. V. Single-cell C(4) photosynthesis versus the dual-cell (Kranz) paradigm. Annu Rev Plant Biol 55, 173–196 (2004). 10.1146/annurev.arplant.55.031903.141725

6 Lundgren, M. R. et al. C4 anatomy can evolve via a single developmental change. Ecology letters 22, 302–312 (2019). 10.1111/ele.13191

7 Hughes, T. E., Sedelnikova, O. V., Wu, H., Becraft, P. W. & Langdale, J. A. Redundant SCARECROW genes patern distinct cell layers in roots and leaves of maize. Development 146 (2019). 10.1242/dev.177543

8 Slewinski, T. L. et al. Short-root1 plays a role in the development of vascular tissue and kranz anatomy in maize leaves. Molecular plant 7, 1388–1392 (2014). 10.1093/mp/ssu036

9 Liu, Q. et al. SHORT ROOT and INDETERMINATE DOMAIN family members govern PIN-FORMED expression to regulate minor vein differentiation in rice. The Plant cell 35, 2848–2870 (2023). 10.1093/plcell/koad125

10 Cribb, L., Hall, L. N. & Langdale, J. A. Four mutant alleles elucidate the role of the G2 protein in the development of C(4) and C(3) photosynthesizing maize tissues. Genetics 159, 787–797 (2001). 10.1093/genetics/159.2.787

11 Ortiz-Ramirez, C. et al. Ground tissue circuitry regulates organ complexity in maize and Setaria. Science 374, 1247–1252 (2021). 10.1126/science.abj2327

12 Ortiz-Ramirez, C., Arevalo, E. D., Xu, X., Jackson, D. P. & Birnbaum, K. D. An Efficient Cell Sorting Protocol forMaize Protoplasts. Current Protocols in Plant Biology 3, e20072 (2018).

13 Bezrutczyk, M. et al. Evidence for phloem loading via the abaxial bundle sheath cells in maize leaves. The Plant cell 33, 531–547 (2021). 10.1093/plcell/koaa055

14 Efroni, I., Ip, P. L., Nawy, T., Mello, A. & Birnbaum, K. D. Quantification of cell identity from single-cell gene expression profiles. Genome biology 16, 9 (2015). 10.1186/s13059-015-0580-x

15 Tian Huang, B. G., Ramin Ranhi, Ken Birnbaum, Doris Wagner A rapid and sensitive multiplex, whole mount RNA fluorescence in situ hybridization and immunohistochemistry protocol. BioRxiv, 2–20 (2023). 10.1101/2023.03.09.531900v1

16 Lundgren, M. R., Osborne, C. P. & Christin, P. A. Deconstructing Kranz anatomy to understand revolution. Journal of experimental botany 65, 3357–3369 (2014). 10.1093/jxb/eru186

17 Chaterjee, J. et al. A low CO2-responsive mutant of Setaria viridis reveals that reduced carbonic anhydrase limits C4 photosynthesis. Journal of experimental botany 72, 3122–3136 (2021). 10.1093/jxb/erab039

18. von Caemmerer, S. Biochemical Models of Leaf Photosynthesis. (CSIRO Pub, 2000).

19 Christin, P. A. & Osborne, C. P. The evolutionary ecology of C4 plants. New Phytol 204, 765–781 (2014). 10.1111/nph.13033

20 Cui, H., Kong, D., Liu, X. & Hao, Y. SCARECROW, SCR-LIKE 23 and SHORT-ROOT control bundle sheath cell fate and function in Arabidopsis thaliana. The Plant journal : for cell and molecular biology 78, 319–327 (2014). 10.1111/tpj.12470

21 Langdale, J. A., Lane, B., Freeling, M. & Nelson, T. Cell lineage analysis of maize bundle sheath and mesophyll cells. Developmental biology 133, 128–139 (1989). 10.1016/0012-1606(89)90304-7

22 Hughes, T. E. & Langdale, J. A. SCARECROW gene function is required for photosynthetic development in maize. Plant Direct 4, e00264 (2020). 10.1002/pld3.264

## Methods references

1 Hao, Y. et al. Dictionary learning for integrative, multimodal and scalable single-cell analysis. Nat Biotechnol (2023).

2 Jin, J. et al. PCMDB: a curated and comprehensive resource of plant cell markers. Nucleic Acids Res 50, D1448–D1455 (2022).

3 Love, M. I., Huber, W. & Anders, S. Moderated estimation of fold change and dispersion for RNA-seq data with DESeq2. Genome biology 15, 550 (2014).

4 Efroni, I., Ip, P. L., Nawy, T., Mello, A. & Birnbaum, K. D. Quantification of cell identity from single-cell gene expression profiles. Genome biology 16, 9 (2015).

5 Birnbaum, K. D. & Kussell, E. Measuring cell identity in noisy biological systems. Nucleic Acids Res 39, 9093–9107 (2011).

6 Huang, T., Guillotin, B., Rahni, R., Birnbaum, K. D. & Wagner, D. A rapid and sensitive, multiplex, whole mount RNA fluorescence in situ hybridization and immunohistochemistry protocol. Plant Methods 19, 131 (2023).

7 Aarón I. Vélez-Ramírez, J. d. D. M., Uriel Pérez-Guerrero, Antonio M. Juarez, Hector Castillo-Arriaga, Josefina Vázquez-Medrano, Ilane Hernández-Morales Open-source LED lamp for the LI-6800 photosynthesis system. bioRxiv, 2–45 (2023).

